# KRT7 Marks an Airway Glandular Ductal Stem Cell Population That Supports Multilineage Development and Regeneration

**DOI:** 10.64898/2026.10.16.759678

**Authors:** Zheqing Hu, Cheng Wang, Yabo Ma, Yaling Yi, Liyuan Yang, Lingling Jiang, Ziying Yan, Xingshen Sun, Xiaoming Liu, John F Engelhardt

## Abstract

Airway submucosal glands (SMGs) play a central role in host defense through mucus secretion, airway hydration, and mucociliary clearance. However, the hierarchical organization of glandular progenitor populations remains poorly defined, largely because SMGs are sparse and restricted to the proximal trachea in murine models. Using the ferret, a model that closely recapitulates the gland-rich submucosal architecture of human cartilaginous airways, we identify KRT7⁺ glandular ductal epithelial cells as a progenitor population. In both adult ferret and healthy human airways, KRT7 expression is primarily restricted to the SMG duct epithelium, where it is co-expressed with KRT19 and CD166 but only partially overlaps with ALDH1A1. During early postnatal development, KRT7 expression becomes progressively restricted to glandular ductal epithelium and is ultimately excluded from mature acinar, myoepithelial cells, and KRT5^+^/KRT14^+^ basal lineages. Fate mapping in *KRT7*^CreER^;*ROSA*^mTmG^ transgenic ferrets revealed that KRT7-lineage cells serve as multipotent progenitors during SMG morphogenesis, generating both serous and mucous epithelial cells. Following severe airway injury, KRT7-lineage cells survive within gland ducts and rapidly expand to repair the damaged surface epithelium by differentiating into ciliated and secretory cells but not KRT5^+^ basal cells. In organoid cultures, glandular KRT7-lineage cells acquired a KRT5⁺ basal-like phenotype and generated multilineage epithelia with functional CFTR-mediated transport. These findings identify the KRT7⁺ ductal niche as a critical progenitor reservoir for airway development, homeostasis, and regeneration, with implications for cystic fibrosis therapies.

## 1 Introduction

Airway submucosal glands (SMGs) are the predominant source of airway secretions in humans and large mammals, with a secretory mass that exceeds that of the surface epithelium ^1,2^. Located within cartilaginous airways, SMGs produce most of the mucus, fluid, and antimicrobial proteins required for airway hydration, mucociliary clearance, and host defense ^3–8^. Their physiological importance is underscored by prominent dysfunction in chronic airway diseases, including chronic obstructive pulmonary disease (COPD) ^9^, asthma ^10,11^, and cystic fibrosis (CF) ^12,13^. In CF, loss of CFTR function impairs chloride and bicarbonate transport, resulting in dehydrated and acidic mucus that obstructs gland ducts from birth and promotes chronic infection and inflammation ^2,3,14–18^. Progressive mucus accumulation and ductal plugging contribute to airway obstruction and declining pulmonary function ^17,19^.

In addition to their secretory functions, SMGs serve as specialized stem cell niches that contribute to airway maintenance and repair ^20–26^. Situated deep within the submucosa, these glandular niches are relatively protected from environmental injury and provide a reservoir of progenitor cells capable of supporting epithelial regeneration. Studies have identified glandular myoepithelial cells (GMECs) and glandular ductal stem cells (GDSCs) as key components of this regenerative hierarchy ^23–27^. Because SMG hypertrophy and hyperplasia are hallmarks of obstructive airway disease, defining the mechanisms that regulate these progenitor populations remains important for the development of regenerative therapies ^23,28,29^.

Current understanding of SMG stem cell plasticity, including the multipotency of GMECs ^25,26^ and GDSCs ^23,24^, derives largely from murine studies. Although these models have provided important insights into glandular stem cell biology, they have limitations for translational lung research ^27,30^. Murine SMGs are sparse and largely restricted to the proximal trachea, and their intrapulmonary airways lack cartilage, as in humans ^31^, limiting investigation of glandular regeneration throughout the conducting airways. Moreover, CF mouse models fail to reproduce the spontaneous glandular pathology and progressive pulmonary disease observed in patients with CF ^32–35^. In contrast, the domestic ferret (*Mustela putorius furo*) closely recapitulates human airway biology ^36–39^. Ferrets possess abundant SMGs throughout cartilaginous airways ^7,40^, and CFTR-knockout ferrets develop spontaneous mucus obstruction, chronic infection, and progressive lung disease that closely resemble human CF ^41–43^.

Among SMG progenitor populations, GDSCs are of particular interest because they contribute to epithelial homeostasis and participate in regeneration following severe injury ^23,44,45^. These cells reside within the gland duct epithelium and are defined by expression of KRT19, CD166 (ALCAM), CD44, and basal cell markers, including KRT5 and KRT14 ^24,44^. GDSCs also exhibit elevated aldehyde dehydrogenase (ALDH) activity, a functional characteristic associated with self-renewal ^46^. Despite the identification of these markers, direct lineage analysis of GDSCs during development and regeneration has remained limited by the absence of genetic tools that selectively label this compartment in vivo.

To address this limitation, we investigated cytokeratin 7 (KRT7) as a candidate marker of the glandular ductal stem cell niche. In the ferret, KRT7 is broadly expressed during lung morphogenesis but becomes progressively restricted to SMG ducts and acini during postnatal maturation ^38,47^. This dynamic spatiotemporal pattern suggested that KRT7 could provide genetic access to a discrete population of glandular progenitors in the adult airway. Establishing KRT7 as a lineage marker would enable direct assessment of GDSC fate, function, and regenerative potential in vivo.

Here, we use a transgenic *KRT7*^CreER^ ferret model to define the developmental and regenerative contributions of KRT7-expressing glandular cells in a species that closely resembles the human airway biology. Through in vivo lineage tracing, we characterize the spatial distribution and developmental restriction of KRT7 expression, determine the contribution of KRT7-lineage cells to SMG morphogenesis, and evaluate their response to epithelial injury. Our findings identify KRT7 as a marker of airway GDSCs and establish this compartment as an important contributor to gland development, maintenance, and regeneration.

## 2 Materials and Methods

### 2.1 Ferret husbandry

All animal procedures were approved by the Institutional Animal Care and Use Committee of the University of Alabama at Birmingham. Male and female domestic ferrets aged 3–24 months were obtained from Marshall Farms (North Rose, NY, USA), whereas neonatal and juvenile ferrets were bred and maintained at the University of Alabama at Birmingham animal facilities. Animals were housed at 20–22°C under a 16-hour light/8-hour dark cycle with ad libitum access to food and water and were acclimated for at least 2 weeks before experimentation. Animals were assigned to experimental groups based on genotype. Investigators were not blinded during experiments or outcome assessment. For studies involving transgenic ferrets, sex was not included as an experimental variable because of limitations in obtaining balanced numbers of male and female animals within each genotype.

### 2.2 Procedure for ferret anesthesia

Animals were anesthetized by subcutaneous injection of ketamine (5-25 mg/kg) and xylazine (0.5-1.5 mg/kg). For intrapulmonary polidocanol administration, anesthesia was maintained with 0.5%–5% isoflurane in oxygen. Throughout the procedures, trained personnel monitored body temperature, pulse rate, oxygen saturation, respiratory rate, and overall clinical condition. Anesthesia was reversed with intramuscular atipamezole (0.5 mg/kg), and animals were monitored until fully recovered before being returned to their home cages.

### 2.3 Lineage tracing in KRT7^CreERT2^;ROSA^mTmG^ ferrets

*KRT7*^CreERT2^ transgenic ferrets were crossed with *ROSA*^mTmG^ Cre-reporter ferrets ^48^ to generate hemizygous offspring. Cre-mediated recombination was induced by intraperitoneal administration of tamoxifen (20 mg/kg in sunflower oil) once daily for 5 consecutive days. Ferrets were euthanized at designated time points for lineage analysis or subjected to polidocanol-induced airway injury 2 weeks after the final tamoxifen dose.

### 2.4 Polidocanol injury of ferret airways

Two weeks after tamoxifen induction, ferrets received a single laryngotracheal administration of polidocanol (2%, 1 mL/kg in saline; Thesit, CAS No. 9002-92-0, Sigma-Aldrich, St. Louis, MO) using a MADgic® Laryngo-Tracheal Mucosal Atomization Device (Teleflex, Morrisville, NC). Animals were monitored daily for activity, body weight, and respiratory rate during the first week after injury and euthanized at designated experimental time points.

### 2.5 Single-molecule fluorescence RNA in situ hybridization

Frozen tracheal and lung sections (10 μm) were analyzed by single-molecule fluorescence RNA in situ hybridization as previously described ^47,49,50^. Ferret *KRT7* probes were designed and synthesized by Thermo Fisher Scientific using ferret accession XM_004774693. Type 1 probes were used to detect ferret *KRT7* mRNA (catalog no. VF6-17245) and human *KRT7* mRNA (assay ID: VA1-10151-VT). Hybridization was performed using the ViewRNA ISH Tissue 2-Plex Assay Kit according to the manufacturer’s instructions (Thermo Fisher Scientific/Affymetrix, Santa Clara, CA). Nuclei were counterstained with DAPI, and images were acquired using a Zeiss LSM880 confocal microscope.

### 2.6 Immunofluorescence staining

For immunofluorescence staining, fixed tissue sections underwent antigen retrieval by boiling in citrate buffer, followed by blocking and permeabilization in PBS containing 10% donkey serum and 0.1% Triton X-100 for 30 min at room temperature. Sections were incubated overnight at 4°C with primary antibodies, washed, and incubated with fluorophore-conjugated secondary antibodies for 1 h at room temperature. Nuclei were counterstained with DAPI, and sections were mounted using ProLong™ Gold Antifade Mountant (Thermo Fisher Scientific). Whole-mount staining was performed as previously described ^25,50^. Briefly, tissues were fixed in 4% paraformaldehyde for 60 min at room temperature, followed by antigen retrieval in sodium citrate buffer with agitation. Samples were blocked and permeabilized in donkey serum containing 0.1% Triton X-100 for 24 h and then incubated sequentially with primary and secondary antibodies for 24 h each at 37°C. After staining, tissues were flattened between glass slides and imaged by confocal microscopy ^51^. Fluorescent images were acquired using Zeiss LSM880 or LSM980 confocal microscopes and processed with Fiji/ImageJ. Antibodies are listed in Supplemental Tables S1 and S2.

### 2.7 Isolation and culture of ferret airway SMG cells

Tracheas and mainstem bronchi were harvested from ferrets, minced with sterile scissors, and transferred to 2-mL screw-cap tubes containing prewarmed pronase solution (1.5 mg/mL). Tissues were incubated at 37°C for 1 h with gentle agitation to remove surface airway epithelial (SAE) cells. Following PBS washes, tissues were digested in 2× collagenase/hyaluronidase buffer (Cat. No. 07912, STEMCELL Technologies) at 37°C for 20 min with continuous shaking. Once the suspension became visibly turbid, 0.25% trypsin-EDTA was added at one-fifth of the total volume, and digestion was continued for an additional 5–10 min to achieve complete dissociation. Enzymatic digestion was terminated by addition of Dulbecco’s Modified Eagle Medium (DMEM) containing 10% fetal bovine serum (FBS). The resulting cell suspension was filtered through a 100-μm cell strainer, centrifuged at 300 × g for 5 min, and resuspended in PneumaCult-Ex Plus airway epithelial culture medium. Cells were seeded onto 804G-coated culture dishes and maintained at 37°C in a humidified incubator with 5% CO₂.

### 2.8 Fluorescence-activated cell sorting

Primary ferret SMG cells were cultured to passage 2, then dissociated using Accutase and incubated at 37°C for 10 minutes to generate single-cell suspensions. Following washing with FACS buffer (PBS supplemented with 2% FBS), cells were filtered through a 40 μm strainer to remove clumps. Fluorescence-activated cell sorting (FACS) was conducted on a BD FACSAria II system. GFP-positive KRT7-lineage cells and tdTomato-positive non-lineage cells were separated via fluorescence. The sorted cells were collected in cold proliferation medium for subsequent experimental procedures.

### 2.9 Polarized epithelium cultured at an air-liquid interface

FACS-sorted GFP⁺ KRT7-lineage and tdTomato⁺ non-lineage cells were expanded to passage 3 in PneumaCult Ex-Plus medium and seeded onto collagen-coated Transwell inserts (0.4-μm pore size, 24-well format) at 2 × 10⁵ cells per insert. Upon reaching confluence, the apical medium was removed to establish an air–liquid interface (ALI), and the basolateral medium was replaced with PneumaCult-ALI differentiation medium. Differentiation medium was refreshed every other day, and cultures were maintained under ALI conditions for 21 days before analysis.

### 2.10 Organoid culture and experimental treatments

FACS-sorted GFP⁺ KRT7-lineage cells and tdTomato⁺ non-lineage cells were mixed at a 1:1 ratio and embedded in growth factor-reduced Matrigel (Cat. No. 354230, Corning). For each well of a 24-well plate, 1,000 GFP⁺ cells and 1,000 tdTomato⁺ cells were suspended in 50 μL Matrigel and plated as domes. After polymerization at 37°C for 15–30 min, cultures were overlaid with 500 μL PneumaCult Ex-Plus medium (STEMCELL Technologies) and maintained for 3 days to allow organoid establishment. On day 3, brightfield and fluorescence images were acquired, and cultures were treated with DMSO vehicle, Alda-1 (MedChemExpress), Daidzin (Sigma-Aldrich; CAS No. 552-66-9), or DEAB (4-diethylaminobenzaldehyde; Sigma-Aldrich; CAS No. 120-21-8). All compounds were used at a final concentration of 20 μM. Treatment medium was refreshed on day 7, and organoids were imaged again on day 10. Cultures were then maintained in PneumaCult ALI medium without drug supplementation until day 21. Organoids were then collected for downstream analyses, including organoid swelling assays and immunostaining. Organoids were fixed, cryoprotected in 20% sucrose, embedded in OCT, cryosectioned, and processed for immunofluorescence staining.

### 2.11 Quantitative analysis of images

All images were analyzed using Fiji/ImageJ. GFP and tdTomato fluorescence channels were processed separately using threshold-based segmentation to identify individual organoids. Organoid number and projected area were measured for each lineage population at the indicated time points, and these data were used to generate quantitative plots of organoid number and size. To assess size distribution, DMSO-treated organoids were categorized into three bins based on projected area: <1,000 μm², 1,000–10,000 μm², and >10,000 μm². The number of GFP-positive and tdTomato-positive organoids in each size bin was quantified for comparative analysis.

### 2.12 Organoid Swelling Assay

To evaluate CFTR-dependent function, day-21 organoids from the DMSO control group were subjected to swelling assays. Organoids were imaged at baseline (0 h) and at the indicated time points following stimulation. Organoid surface area was measured using Fiji/ImageJ and normalized to the baseline value for each organoid. Changes in area were expressed as a percentage increase relative to baseline ^52–54^. The integrated area under the curve (iAUC) was calculated to quantify overall swelling responses ^53,54^.

### 2.13 Ussing chamber short circuit current (Isc) assays

Transepithelial ion transport was measured using an Ussing chamber system (U2500, Physiologic Instruments) equipped with voltage-clamp and perfusion controls. ALI cultures grown on Transwell inserts were mounted between hemichambers containing warmed (37°C), oxygenated Ringer’s solution continuously gassed with 95% O₂/5% CO₂. Chloride currents were measured under asymmetric Cl⁻ conditions. The basolateral chamber contained high-Cl⁻ Ringer’s solution (135 mM NaCl, 1.2 mM CaCl_2_, 1.2 mM MgCl_2_, 2.4 mM KH_2_PO_4_, 0.2 mM K_2_HPO_4_, 5 mM HEPES; pH 7.4), whereas the apical chamber contained low-Cl⁻ solution in which NaCl was replaced with 135 mM sodium gluconate. Epithelia were voltage-clamped at 0 mV and short-circuit current (Isc) was recorded using Ag/AgCl electrodes connected by 3 M KCl agar bridges as previously described ^54,55^.

To assess CFTR-mediated Cl⁻ secretion, drugs were added sequentially to the apical chamber: amiloride (10 μM) to inhibit ENaC-mediated Na⁺ absorption, DIDS (50 μM) to inhibit non-CFTR chloride channels, forskolin (10 μM) and IBMX (100 μM) to elevate intracellular cAMP and activate CFTR, and GlyH101 (50 μM) to inhibit CFTR. Bumetanide (100 μM) was added basolaterally to inhibit NKCC-dependent chloride uptake. Changes in short-circuit current (ΔIsc) were calculated from steady-state currents before and after each treatment. Data were acquired using Acquire & Analyze software (Physiologic Instruments) and analyzed in GraphPad Prism.

### 2.14 Statistical analyses

Statistical analyses were performed using GraphPad Prism v9.0 (GraphPad Software). Comparisons among multiple groups were analyzed by one-way ANOVA followed by Dunnett’s multiple-comparison test. Comparisons between two groups were performed using paired or unpaired Student’s t-tests, as appropriate. Data are presented as mean ± SEM unless otherwise indicated. Differences were considered statistically significant at P < 0.05.

## 3 Results

### 3.1 KRT7 Marks a Conserved Glandular Ductal Compartment in Ferret and Human Airways

Unlike mice, which possess sparse SMGs largely restricted to the proximal trachea, humans and ferrets contain abundant SMGs throughout the cartilaginous airways ^45,56^. Alcian Blue staining revealed similar gland-rich submucosal architecture in ferret and human bronchi, characterized by abundant SMGs connected to the airway lumen through well-defined gland ducts (GDs) (Suppl. Fig. S1A,B). In contrast, murine airways exhibited a markedly reduced submucosal compartment, with sparse SMGs confined to the proximal trachea (Suppl. Fig. S1C). These findings support the ferret as a physiologically relevant model for studies of airway gland biology and mucus-associated airway disease.

Because the gland duct compartment serves as a niche for glandular ductal stem cells (GDSCs) ^23,24^, we sought to identify markers that define this population. KRT7 is a well-established marker of glandular duct epithelium across multiple organs ^57–59^. Because KRT7 marks epithelial populations with progenitor activity in other tissues ^60^, we examined its distribution in ferret and human airways. Single-molecule RNA in situ hybridization demonstrated that KRT7 transcripts were largely restricted to the gland duct epithelium in both ferret trachea and human bronchi, with minimal expression in the surface airway epithelium (SAE) or deep acinar compartments (Suppl. Fig. S2A–D).

To determine whether KRT7-expressing cells reside within the GDSC niche, we examined their relationship to established ductal stem cell markers. In both species, KRT7-positive cells co-localized with KRT19, CD166, and ALDH1A1 (Fig. 1A–F) ^44,46^. KRT7 exhibited extensive overlap with KRT19 and CD166 throughout the ductal epithelium, while overlap with ALDH1A1 was more regionally restricted, suggesting heterogeneity within the ductal compartment. The highly similar expression patterns observed in ferret and human airways identify KRT7 as a conserved marker of the glandular ductal stem cell niche and further support the ferret as a relevant model for human airway gland biology.

**Figure 1.**
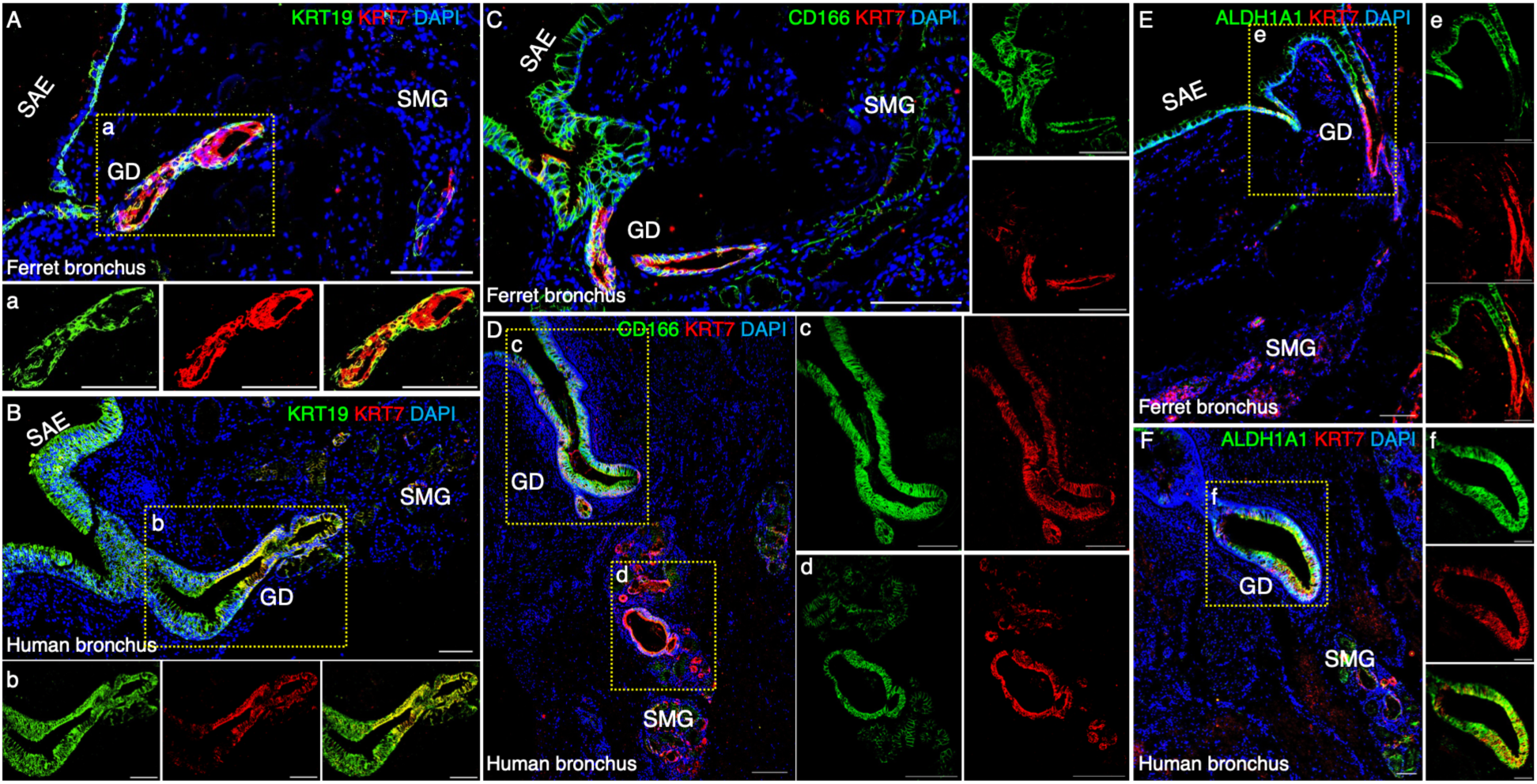
Conserved expression of KRT7 with established glandular ductal stem cell (GDSC) markers in ferret and human airway submucosal glands. Immunofluorescence staining of healthy adult ferret and human bronchial sections was performed to examine the spatial distribution of KRT7 relative to the established GDSC markers KRT19, CD166, and ALDH1A1. (A–F) Representative images showing KRT7 (red) together with KRT19 (green; A, B), CD166 (green; C, D), or ALDH1A1 (green; E, F) in bronchi from ferrets (A, C, E) and humans (B, D, F). Bottom panels (a, b) and right-side panels (c–f) show higher-magnification views of the boxed regions in the corresponding main images, including merged and single-channel fluorescence images. KRT7 extensively co-localizes with KRT19 and CD166 within the gland duct (GD) epithelium of submucosal glands (SMGs), whereas overlap with ALDH1A1 is limited to a subset of KRT7-expressing cells, indicating heterogeneity within the ductal progenitor compartment. Nuclei were counterstained with DAPI. Scale bars, 100 μm. GD, gland duct; SAE, surface airway epithelium; SMG, submucosal gland.

### 3.2 KRT7 Expression Progressively Restricts to Adult Glandular Ductal Cells During Development

Ferret SMGs undergo extensive postnatal development during the first several weeks of life ^40,61^. To determine how the adult KRT7^+^ GD compartment is established, we examined KRT7 expression during SMG maturation from birth to six months of age (Fig. 2; Suppl. Fig. S3). During early postnatal development, KRT7 was broadly expressed throughout the SAE and invaginating gland buds (Fig. 2A–C), where it co-localized with the gland morphogenesis regulator LEF1 ^62^. As gland development progressed, KRT7 expression became increasingly restricted to the developing GD epithelium (Fig. 2D–G). Concurrently, KRT7 segregated from differentiating Lysozyme^+^ serous cells, MUC5B^+^ mucous cells, and α-smooth muscle actin (SMA) positive glandular myoepithelial cells (GMECs) ^25–27^. By six months of age, KRT7 expression was primarily confined to the GD epithelium and excluded from mature acinar and myoepithelial compartments in both tracheal and bronchial airways (Fig. 2H,I). A similar pattern was observed relative to basal cell markers (Suppl. Fig. S3A–D). At birth, KRT7 was broadly co-expressed with KRT5 and KRT14 within primitive gland buds. As development progressed, these populations diverged, and by adulthood, KRT5 and KRT14 marked basal and myoepithelial populations, whereas KRT7 remained restricted to the GD epithelium. These findings demonstrate that KRT7 transitions from a broadly expressed epithelial marker during early morphogenesis to a lineage-restricted marker of the adult ductal compartment.

**Figure 2.**
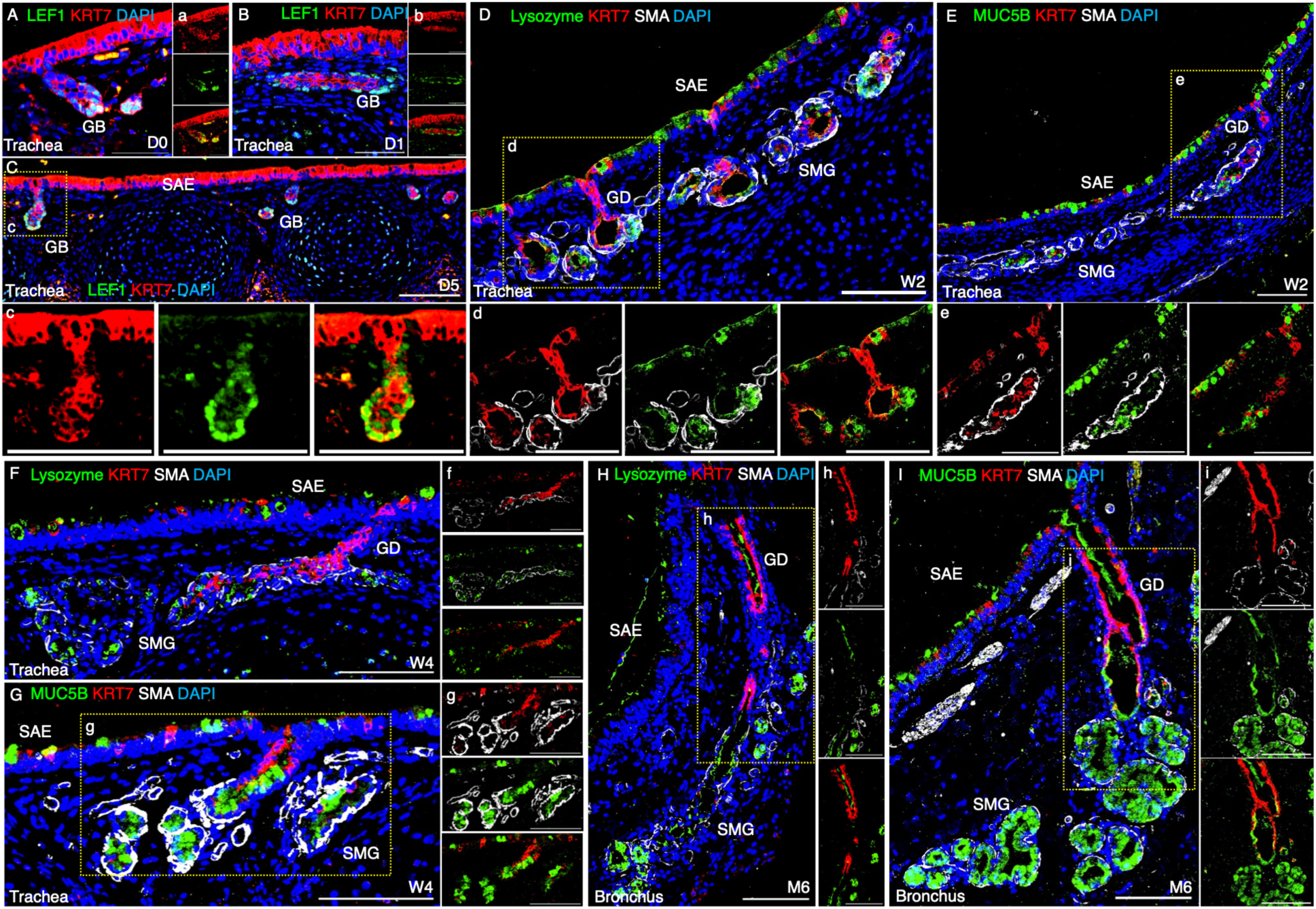
Spatiotemporal restriction of KRT7 to the gland duct lineage during ferret airway submucosal gland (SMG) development. Immunofluorescence staining was performed on ferret tracheal and bronchial sections collected during postnatal development [Day 0 (D0), Day 1 (D1), Day 5 (D5), Week 2 (W2), Week 4 (W4), and Month 6 (M6)] to define the spatial distribution of KRT7 relative to lineage-specific markers. (A–C) Representative images of D0–D5 trachea showing KRT7 (red) and lymphoid enhancer-binding factor 1 (LEF1; green), a transcription factor marking early SMG morphogenesis. KRT7 and LEF1 are co-expressed within invaginating epithelial gland buds, while KRT7 is broadly expressed throughout the surface airway epithelium (SAE). (D–G) Representative W2–W4 tracheal sections stained for KRT7 (red), α-smooth muscle actin (SMA; gray), and either the serous cell marker lysozyme (green; D, F) or the mucous cell marker MUC5B (green; E, G). During gland maturation, KRT7 expression becomes progressively restricted to the developing gland ducts and is excluded from differentiating serous (lysozyme⁺), mucous (MUC5B⁺), and myoepithelial (SMA⁺) lineages. (H, I) Representative adult (M6) bronchial sections demonstrate persistent localization of KRT7 to the stratified gland duct epithelium, with no detectable expression in mature serous, mucous, or myoepithelial cells. Dashed boxes indicate regions shown at higher magnification in the corresponding lowercase panels (c–i), which include merged and single-channel images. Nuclei were counterstained with DAPI. Scale bars, 100 μm. GB, gland bud; GD, gland duct; SAE, surface airway epithelium; SMG, submucosal gland.

To determine whether KRT7 labels a stable adult ductal population, we performed short-term lineage tracing in tamoxifen-induced *KRT7*^CreER^;*ROSA*^mTmG^ ferrets (Fig. 3A) (Yi et al., submitted manuscript). Two weeks after induction, GFP labeling was predominantly confined to gland ducts in both tracheal and bronchial airways, whereas the SAE and acinar compartments remained tdTomato^+^, with only rare GFP^+^ cells observed in acini adjacent to gland ducts (Suppl. Fig. S4A,B). Within the ductal compartment, GFP expression overlapped with endogenous KRT7, indicating stable labeling of resident KRT7-expressing cells rather than a transient developmental population (Fig. 3B–D). To further define these lineage-labeled cells, we examined established markers of adult glandular ductal stem cells (GDSCs). GFP^+^ cells co-expressed KRT19, CD166, and ALDH1A1 within the GD epithelium (Fig. 3C–E) ^24,44,46^, with high-magnification images confirming triple-positive cells within individual ducts. Similar duct-restricted labeling was observed in esophageal SMGs, whereas salivary glands exhibited only sparse GFP labeling (Suppl. Fig. S4C,D). Together, these findings identify KRT7 as a stable marker of a resident ductal epithelial population that overlaps with previously defined GDSC niches in adult ferret airways.

**Figure 3.**
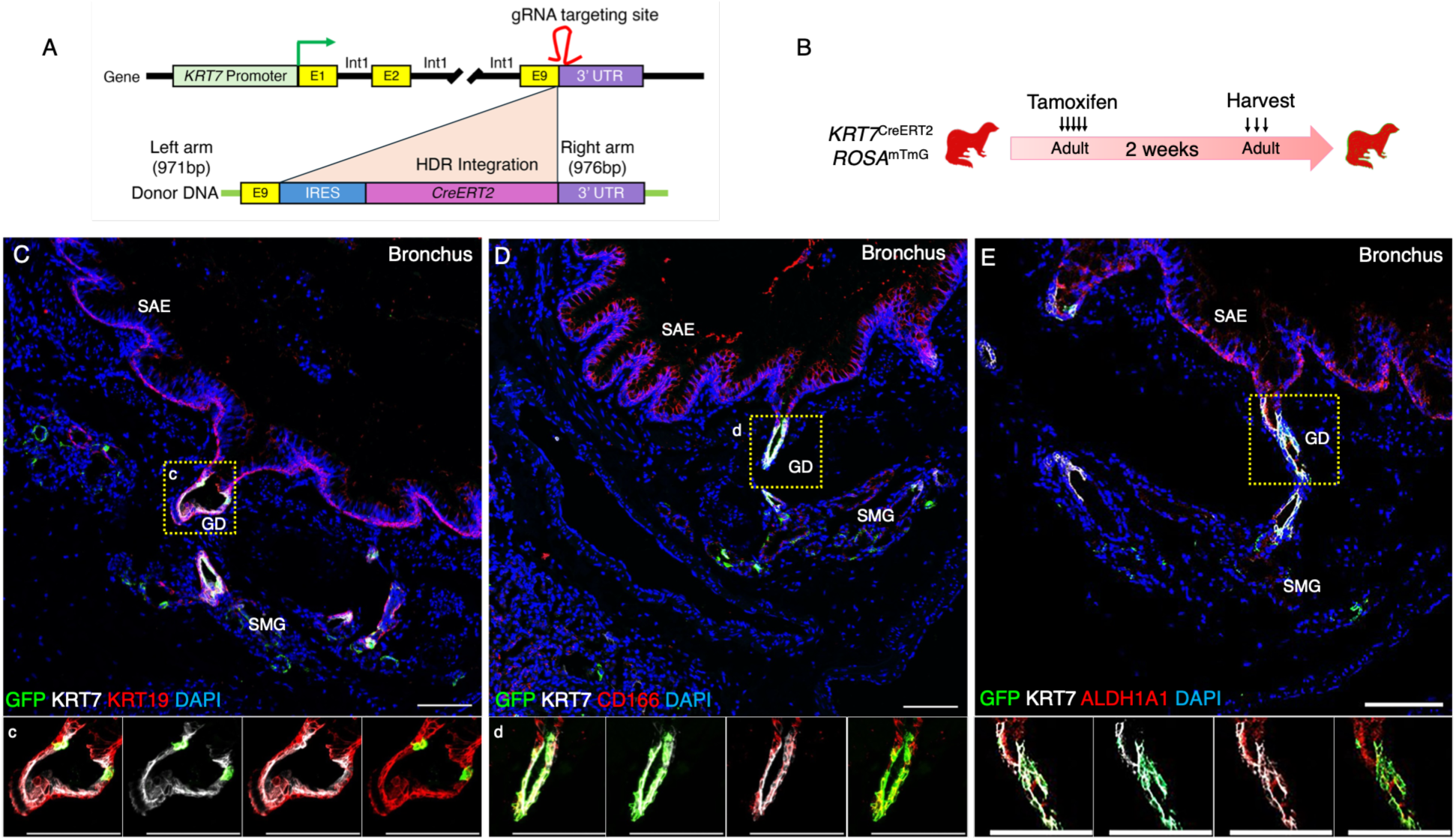
KRT7 marks resident subpopulations of glandular ductal stem cells in adult ferret airways. (A) Schematic illustrating generation of the KRT7-IRES-CreERT2 (*KRT7*^CreER^) ferret. An IRES-CreERT2 cassette was inserted into the 3′ untranslated region (UTR) immediately downstream of the endogenous *KRT7* stop codon by CRISPR/Cas9-mediated homology-directed repair (HDR). Breeding *KRT7*^CreER^ ferrets with the *ROSA*^mTmG^ reporter line generated *KRT7*^CreER^;*ROSA*^mTmG^ lineage-tracing ferrets. (B) Experimental design for short-term lineage labeling. Adult *KRT7*^CreERT2^;*ROSA*^mTmG^ ferrets received tamoxifen, and airway tissues were collected 2 weeks later to examine co-expression of GFP-labeled KRT7-lineage cells with established glandular ductal stem cell (GDSC) markers. (C–E) Representative immunofluorescence images of adult bronchial sections showing GFP-positive KRT7-lineage cells (green), endogenous KRT7 protein (white), and KRT19 (red; C), CD166 (red; D), or ALDH1A1 (red; E). GFP-positive lineage cells largely retain KRT7 expression and co-express KRT19 and CD166, whereas overlap with ALDH1A1 is limited to a subset of cells, consistent with heterogeneity within the gland duct compartment. Dashed boxes indicate regions shown at higher magnification in the corresponding lowercase panels (c–e), which include merged and double-channel images. Nuclei were counterstained with DAPI. Scale bars, 100 μm. GD, gland duct; SAE, surface airway epithelium; SMG, submucosal gland.

### 3.3 Glandular KRT7^+^ Cells Generate Serous and Mucous Lineages During Postnatal SMG Development

To determine the developmental potential of KRT7-expressing cells, we performed lineage tracing during active postnatal SMG morphogenesis in *KRT7*^CreER^;*ROSA*^mTmG^ ferrets. Tamoxifen was administered for five consecutive days beginning at 2 weeks of age (W2), and tissues were collected at 4 weeks (W4) and 6 months (M6) (Fig. 4A). At W4, GFP^+^ KRT7-lineage cells were primarily confined to the ductal epithelium with extension into adjacent glandular structures (Fig. 4B). By M6, lineage-labeled cells still occupied extensive regions of the SMG network while remaining connected to their ductal origin (Fig. 4C). Although KRT7 protein expression remained restricted to gland ducts, GFP^+^ descendants populated differentiated glandular compartments, indicating loss of KRT7 expression during lineage maturation. KRT7-lineage cells contributed extensively to both Lysozyme^+^ serous and MUC5B^+^ mucous cell populations (Fig. 4D,E,H,I). In contrast, GFP^+^ cells were not detected in MUC5AC^+^ surface goblet cells, FOXJ1^+^ or acetylated α-tubulin^+^ ciliated cells at either time point (Fig. 4C,F–I). Morphometric analysis demonstrated a stable proportion of KRT7-lineage descendants during SMG maturation from W4 and M6 (Fig. 4J–L). Notably, more GFP^+^/KRT5^+^ basal cells were more abundant in the GD epithelium at W4 than at M6 (Fig. 4F,G,K,L), suggesting that KRT7-lineage cells transiently contribute to the ductal basal cell population during gland maturation. These findings demonstrate that postnatal KRT7^+^ cells function as gland-restricted progenitors that generate serous and mucous SMG lineages while excluding surface epithelial and ciliated cell fates during gland development.

**Figure 4.**
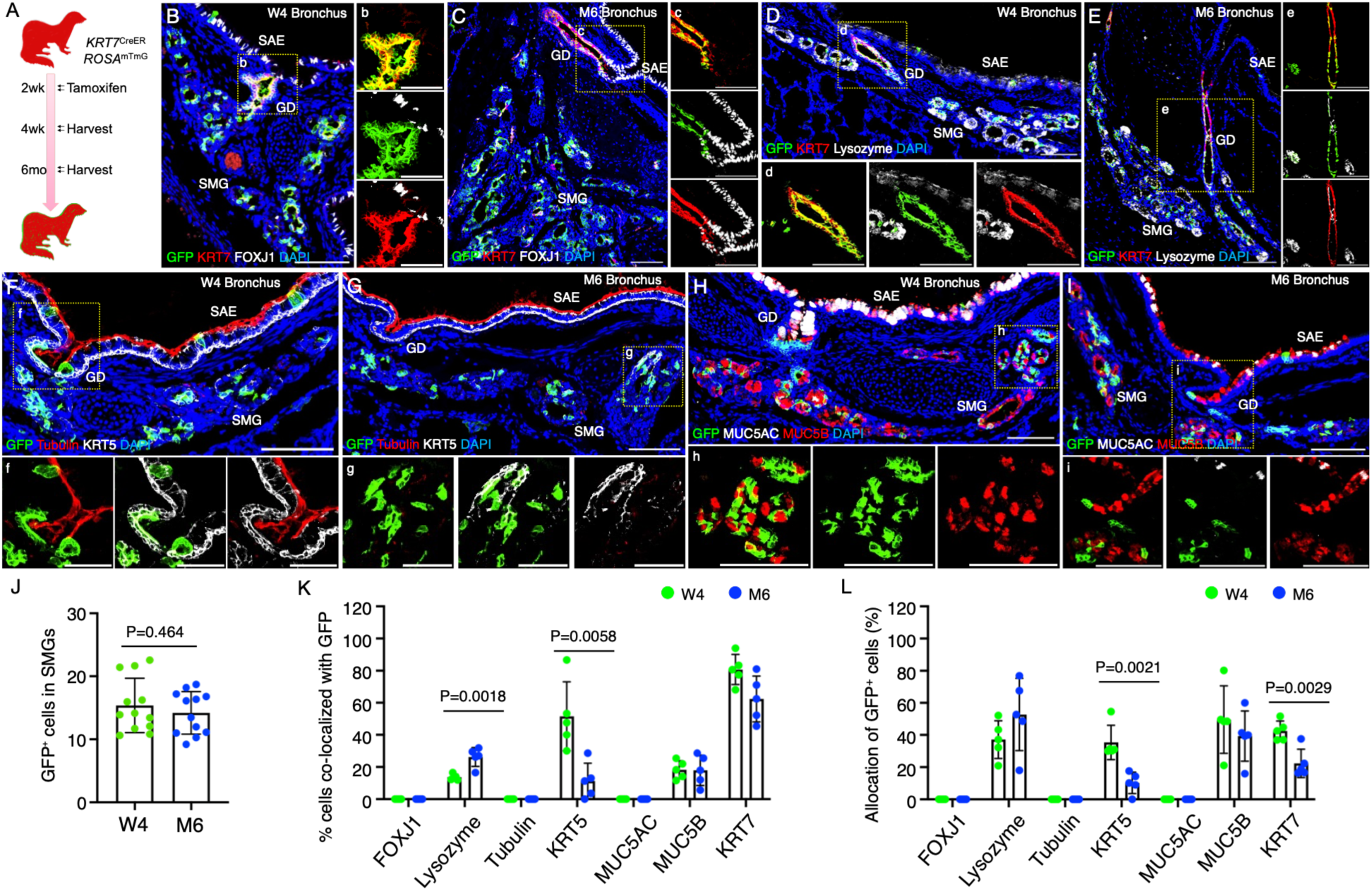
KRT7-lineage cells function as multipotent progenitors during postnatal airway submucosal gland (SMG) development in the ferret. (A) Experimental design for postnatal KRT7 lineage tracing. *KRT7*^CreER^;*ROSA*^mTmG^ ferrets received tamoxifen for five consecutive days beginning at 2 weeks of age. Airway tissues were collected at 4 weeks (W4), during active SMG development, and at 6 months (M6), after gland maturation. (B, C) Representative immunofluorescence images of W4 (B) and M6 (C) bronchi stained for GFP (green), KRT7 (red), and the ciliated cell marker FOXJ1 (gray). GFP-positive KRT7-lineage cells expand from the gland duct (GD) into the developing SMG but do not contribute to FOXJ1-positive ciliated cells. (D–I) Representative W4 (D, F, H) and M6 (E, G, I) bronchial sections demonstrating multilineage differentiation of KRT7-lineage cells. GFP-positive descendants co-localize extensively with the serous cell marker lysozyme (gray; D, E) and the mucous cell marker MUC5B (red; H, I), indicating differentiation into both serous and mucous glandular cells. In contrast, GFP does not co-localize with KRT5-positive basal cells (gray; F, G), acetylated α-tubulin-positive ciliated cells (red; F, G), or MUC5AC-positive surface goblet cells (gray; H, I). Nuclei were counterstained with DAPI. Lowercase panels show higher-magnification views or individual fluorescence channels of the corresponding boxed regions. (J–L) Quantification of KRT7-lineage tracing. (J) Percentage of GFP-positive cells within the SMG (including the gland duct) at W4 and M6. (K) Percentage of each epithelial cell population that was GFP-positive (GFP/marker double-positive cells within each lineage). (L) Distribution of GFP-positive lineage cells among glandular cell types (GFP/marker double-positive cells as a percentage of total GFP-positive glandular cells). Data are presented as mean ± SEM. Statistical significance was determined using an unpaired Welch’s t-test (J) and multiple unpaired t-tests (K, L). Individual data points represent biological replicates. Scale bars, 100 μm (B, F, G) and 50 μm (C–E, H, I). GD, gland duct; SAE, surface airway epithelium; SMG, submucosal gland.

### 3.4 KRT7-Lineage Cells Contribute to Surface Airway Epithelial Regeneration Following Severe Injury

Previous studies in mice have shown that GDSCs and GMECs survive severe airway injury and contribute to epithelial repair ^24–26,44,63^. To determine whether glandular KRT7-lineage cells possess similar regenerative capacity in the ferret airway, adult *KRT7*^CreER^;*ROSA*^mTmG^ ferrets were subjected to polidocanol-induced injury two weeks after tamoxifen induction (Fig. 5A). Tissues were analyzed at 2 and 14 days post-injury (DPI). At 2 DPI, the SAE was extensively depleted, whereas GFP^+^ KRT7-lineage cells persisted within GDs (Fig. 5B,D). Surviving lineage-labeled cells retained KRT7 expression within the GD epithelium (Fig. 5E) but did not co-localize with KRT5 at either 2 or 14 DPI, indicating little contribution to the basal cell compartment (Fig. 5H,I). By 14 DPI, GFP^+^ KRT7-lineage cells had expanded markedly and migrated from the GDs onto the repairing airway surface (Fig. 5B,D). Consistent with activation of a regenerative program, SOX9 expression was detected in cells emerging from the GDs during repair (Fig. 5C). A subset of lineage-labeled cells on the regenerating surface retained KRT7 expression (Fig. 5F).

**Figure 5.**
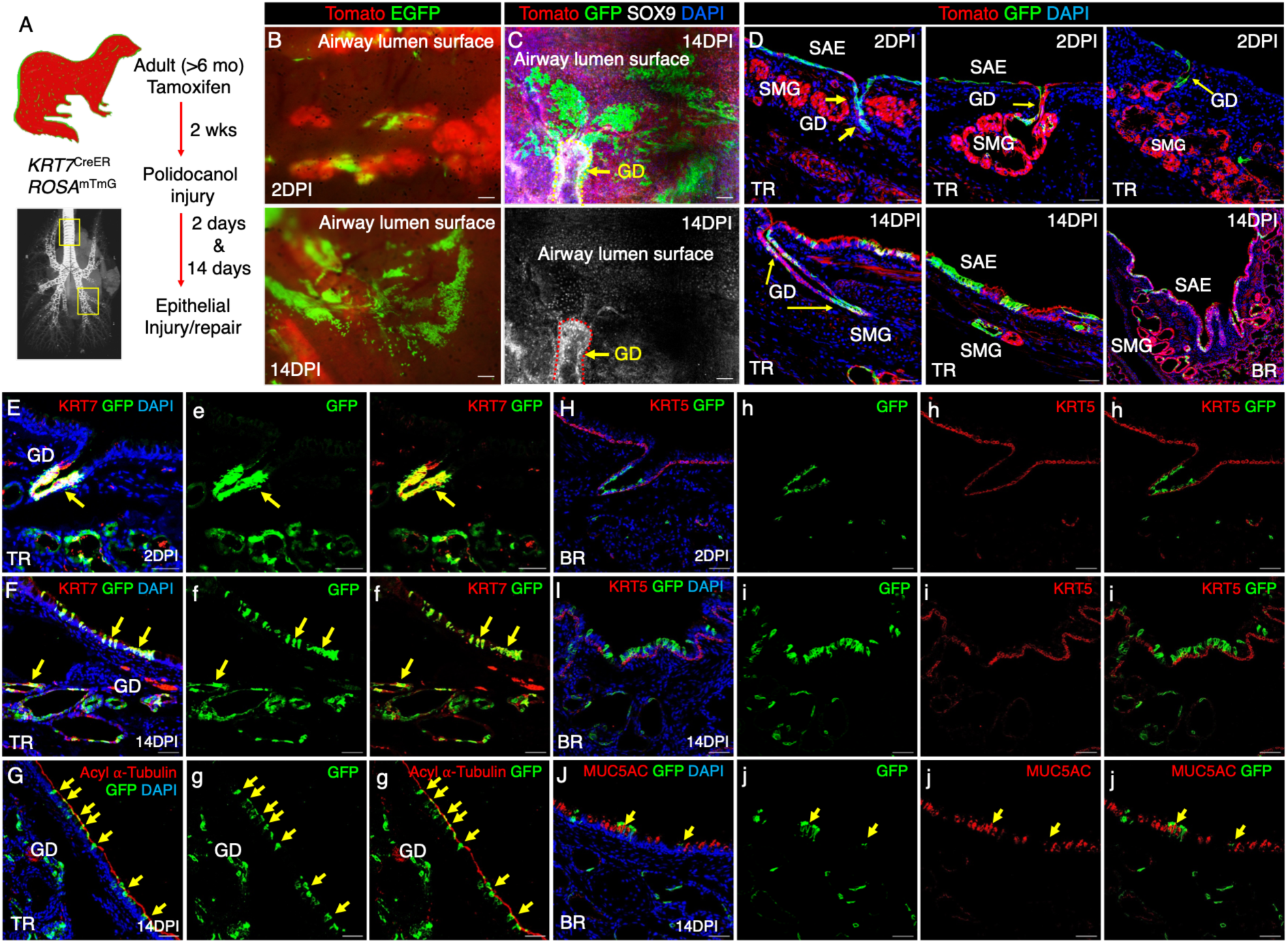
KRT7-lineage cells contribute to multilineage epithelial regeneration following severe surface airway injury. (A) Experimental design. Adult *KRT7*^CreER^;*ROSA*^mTmG^ ferrets underwent lineage labeling with tamoxifen followed by intratracheal polidocanol injury 2 weeks later. Airway tissues were collected at 2 days post-injury (2 DPI) and 14 days post-injury (14 DPI) to evaluate epithelial regeneration. (B) Representative whole-mount fluorescence images of the tracheal surface at 2 DPI and 14 DPI. GFP-positive KRT7-lineage cells initially appear as small focal patches at 2 DPI and expand extensively across the regenerating airway surface by 14 DPI. (C) Representative whole-mount immunofluorescence image at 14 DPI showing SOX9 (white) expression in GFP-positive cells emerging from the gland duct (GD) onto the regenerating airway surface. The lower panel shows the SOX9 channel alone. (D) Representative tracheal (TR) and bronchial (BR) sections at 2 DPI (top) and 14 DPI (bottom). At 2 DPI, GFP-positive KRT7-lineage cells remain largely confined to the gland ducts. By 14 DPI, GFP-positive cells have expanded from the ducts into the regenerating surface airway epithelium (SAE). Unrecombined cells remain tdTomato-positive (red). (E–J) Immunofluorescence analysis of lineage differentiation during epithelial repair. GFP-positive lineage cells co-express KRT7 within the gland duct at 2 DPI (E) and within both the gland duct and regenerated SAE at 14 DPI (F). At 14 DPI, GFP-positive cells differentiate into acetylated α-tubulin-positive ciliated cells (G) and MUC5AC-positive goblet cells (J), but do not contribute to KRT5-positive basal cells at either 2 DPI (H) or 14 DPI (I). Arrows indicate GFP-positive lineage cells. Lowercase panels show higher-magnification views or individual fluorescence channels corresponding to the boxed regions. Nuclei were counterstained with DAPI. Scale bars, 100 μm (B, C) and 50 μm (D–J). BR, bronchus; DPI, days post-injury; GD, gland duct; SAE, surface airway epithelium; SMG, submucosal gland; TR, trachea.

To determine whether KRT7-lineage cells generated differentiated epithelial populations, we examined lineage-specific markers in the regenerated SAE. At 14 DPI, GFP^+^ cells co-localized with acetylated α-tubulin and MUC5AC, indicating differentiation into ciliated and goblet cell lineages, respectively (Fig. 5G,J). In contrast, GFP^+^ cells remained absent from the KRT5^+^ basal compartment (Fig. 5I). Together, these findings demonstrate that KRT7-lineage ductal cells function as reserve progenitors that survive severe injury, expand from the glandular duct compartment, and contribute directly to the regeneration of the luminal airway epithelium.

### 3.5 KRT7-Lineage Cells Exhibit Enhanced Clonogenic Growth and ALDH-Responsive Expansion

Because elevated aldehyde dehydrogenase (ALDH) activity enriches airway progenitor populations with enhanced self-renewal capacity ^46^, we examined the proliferative potential of KRT7-lineage cells isolated from SMGs of tamoxifen-induced *KRT7*^CreERT2^;*ROSA*^mTmG^ ferrets (Suppl. Fig. S5). Following dissociation, GFP^+^ KRT7-lineage cells and tdTomato^+^ non-lineage cells were cultured on plastic prior to fluorescence-activated cell sorting (FACS) (Suppl. Fig. S5). By day 7, GFP^+^ KRT7-lineage cells formed large cobblestone colonies, consistent with robust proliferative capacity (Suppl. Fig. S5A). FACS yielded highly purified and viable GFP^+^ KRT7-lineage cells (32.72 ± 7.02%) and tdTomato+ non-lineage cells (62.62 ± 7.81%) populations for downstream functional analyses (Suppl. Fig. S5B–D).

To compare clonogenic potential, GFP^+^ KRT7-lineage cells and tdTomato^+^ non-lineage cells were co-cultured at a 1:1 ratio in 3D organoid assays under proliferative conditions (Fig. 6A). After 3 days, both populations contributed similarly to early organoid structures, indicating comparable initial seeding and survival. In contrast, after 10 days, a subset of GFP^+^ KRT7-lineage organoids were larger than tdTomato^+^ controls under vehicle-treated conditions (Fig. 6B). Quantitative analysis demonstrated that the largest organoids (>10,000 μm²) were preferentially derived from the GFP^+^ fraction, whereas smaller organoids were represented in both populations (Fig. 6C). These findings indicate that KRT7-lineage cells possess enhanced proliferative capacity while maintaining similar clonogenic potential to non-lineage cells.

**Figure 6.**
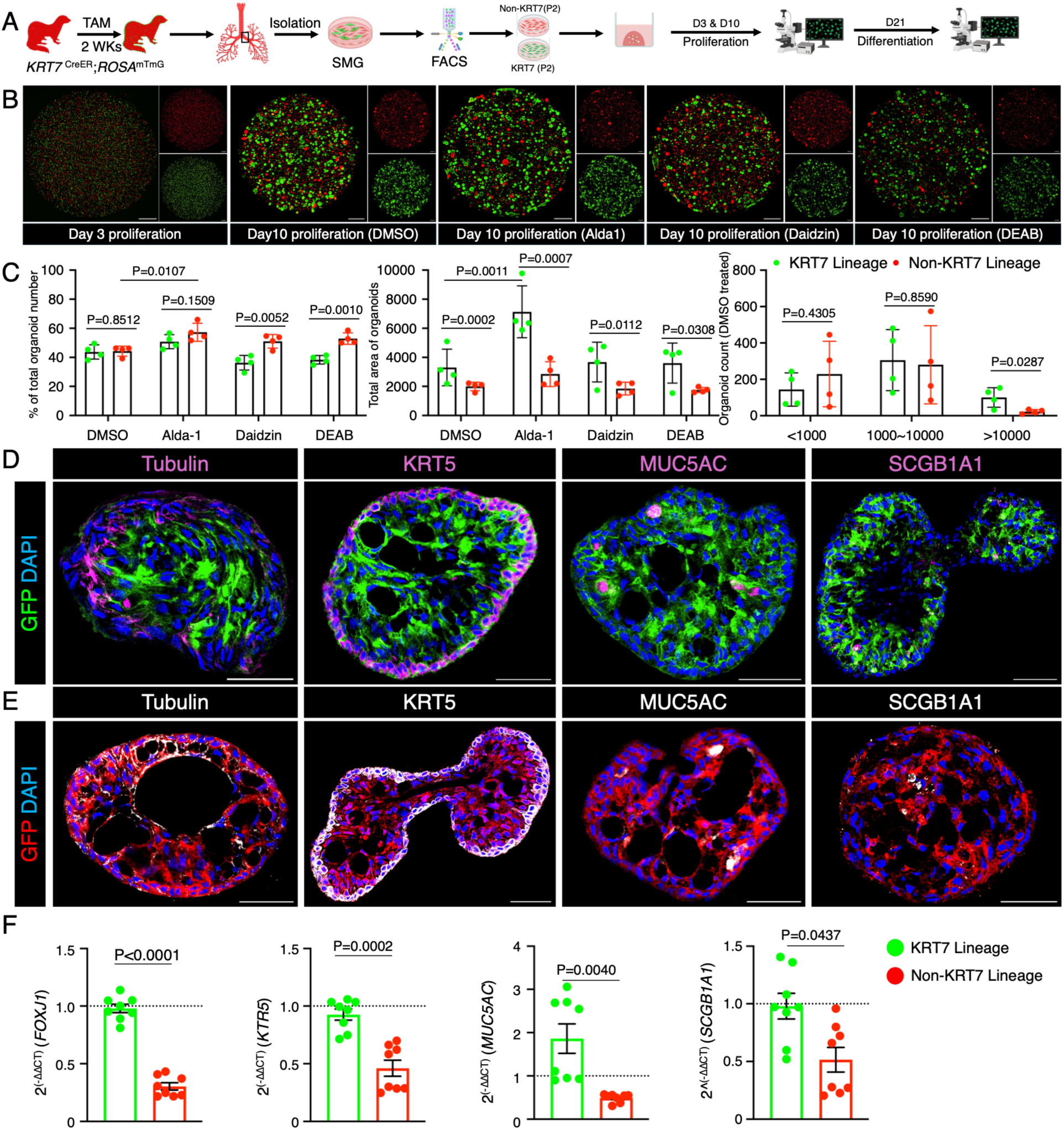
KRT7-lineage glandular ductal cells exhibit enhanced proliferative capacity and multilineage differentiation in organoid culture. (A) Experimental design. Adult *KRT7*^CreER^;*ROSA*^mTmG^ ferrets received tamoxifen for lineage labeling before SMG isolation and fluorescence-activated cell sorting (FACS) of GFP-positive KRT7-lineage and tdTomato-positive non-lineage cells. Sorted cells were mixed at a 1:1 ratio and cultured as organoids. From day 3 (D3) to day 10 (D10), organoids were maintained in PneumaCult Ex-Plus medium containing DMSO, Alda-1 (ALDH activator), Daidzin (ALDH2 inhibitor), or DEAB (pan-ALDH inhibitor). Cultures were then switched to PneumaCult ALI medium until day 21 (D21) for differentiation. (B) Representative whole-well fluorescence images of organoids at D3 and D10 under the indicated treatment conditions. GFP-positive KRT7-lineage organoids (green) and tdTomato-positive non-lineage organoids (red) are shown together, with corresponding single-channel images displayed at right. (C) Quantification of organoid growth at D10. Left, proportion of total organoids derived from GFP-positive and tdTomato-positive cells. Middle, total organoid area per well. Right, size distribution of organoids under DMSO control conditions (<1,000, 1,000–10,000, and >10,000 μm²). Data are from N = 4 independent cultures generated from two donor ferrets. (D, E) Representative immunofluorescence images of D21 organoids derived from GFP-positive KRT7-lineage cells (D) or tdTomato-positive non-lineage cells (E), stained for acetylated α-tubulin (ciliated cells), KRT5 (basal cells), MUC5AC (goblet cells), and SCGB1A1 (club cells), demonstrating multilineage differentiation. (F) TaqMan RT-PCR analysis of *FOXJ1*, *KRT5*, *MUC5AC*, and *SCGB1A1* expression in GFP-positive and tdTomato-positive organoids at D21 (N = 8 independent cultures from four experiments using two donor ferrets). Nuclei were counterstained with DAPI. Scale bars, 500 μm (B) and 50 μm (D, E). Data are presented as mean ± SEM. Statistical significance was determined using multiple unpaired t-tests (C) and paired t-tests (F). Individual data points represent biological replicates.

To determine whether this growth advantage was associated with ALDH activity, organoids were treated with pharmacologic modulators of the ALDH pathway. ALDH activation with Alda-1 further enhanced organoid growth, whereas ALDH inhibition with Daidzin or DEAB attenuated expansion of KRT7-lineage organoids (Fig. 6B,D). Alda-1 significantly increased organoid size in the KRT7-lineage population without altering organoid number, while increasing the number of organoids derived from the non-lineage fraction. In contrast, ALDH inhibition had minimal effects on organoid number or size in either population. Together, these findings indicate that KRT7-lineage cells possess an intrinsic proliferative advantage that is modulated by ALDH signaling.

### 3.6 KRT7-Lineage Cells Generate Multilineage Airway Epithelia with Functional CFTR Activity

To evaluate differentiation potential, organoids derived from KRT7-lineage and non-lineage cells were cultured under epithelial differentiation conditions. Both populations generated acetylated α-tubulin^+^ ciliated cells, KRT5^+^ basal cells, MUC5AC^+^ goblet cells, and SCGB1A1^+^ secretory (Fig. 6E,F); however, KRT7-lineage organoids demonstrated increased expression of *FOXJ1* (ciliated cell marker), *KRT5*, *MUC5AC*, and *SCGB1A1* relative to non-lineage controls (Fig. 6G). Similar results were observed under air-liquid interface (ALI) conditions, where KRT7-lineage cultures generated more extensively differentiated epithelia at both the protein and transcript levels (Fig. 7A–C).

**Figure 7.**
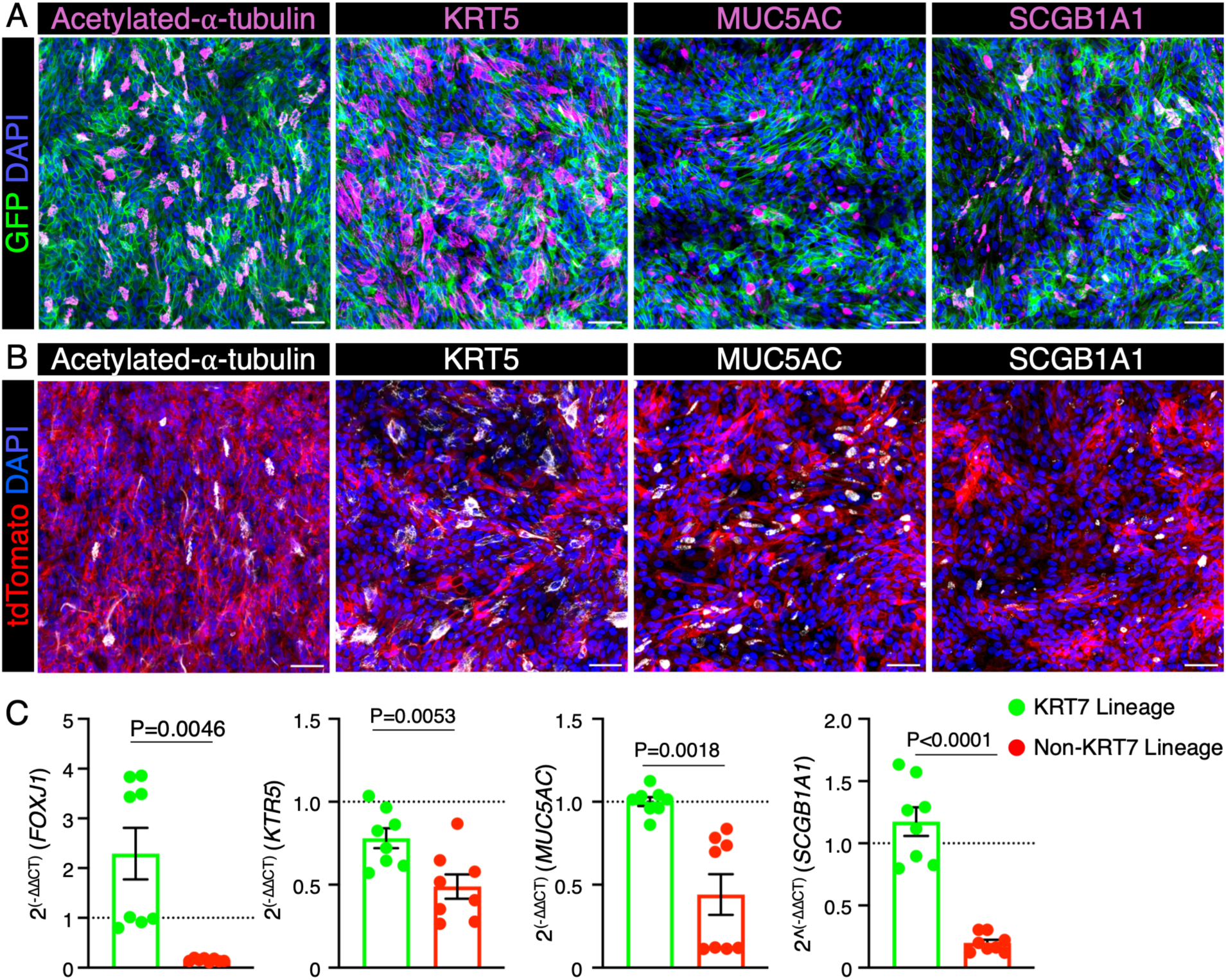
KRT7-lineage glandular ductal cells exhibit enhanced multilineage differentiation in polarized air-liquid interface (ALI) cultures. (A, B) Representative immunofluorescence images of differentiated ALI cultures derived from GFP-positive KRT7-lineage cells (A) and tdTomato-positive non-lineage cells (B), stained for acetylated α-tubulin (ciliated cells), KRT5 (basal cells), MUC5AC (goblet cells), and SCGB1A1 (club cells). (C) TaqMan RT-PCR analysis of *FOXJ1*, *KRT5*, *MUC5AC*, and *SCGB1A1* expression in differentiated ALI cultures derived from GFP-positive KRT7-lineage and tdTomato-positive non-lineage cells (N = 8 independent cultures from four experiments using two donor ferrets). Data are presented as mean ± SEM. Statistical significance was determined using an unpaired t-test. Individual data points represent biological replicates. Nuclei were counterstained with DAPI. Scale bars, 20 μm.

To assess epithelial function, we examined CFTR-dependent fluid and ion transport (Suppl. Fig. S6A). Forskolin stimulation induced organoid swelling in KRT7-lineage cultures that was significantly greater than that observed in non-lineage controls (P < 0.0001) (Suppl. Fig. S6A–D). Such forskolin-induced swelling activity has typically been associated with CFTR function ^54^. Short-circuit current measurements further confirmed CFTR-mediated ion transport in both lineage-and non-lineage-derived ALI cultures that was not significantly different (Suppl. Fig. S6E,F). Together, these findings demonstrate that KRT7-lineage cells generate multilineage airway epithelia capable of CFTR-mediated fluid and ion transport.

## 4 Discussion

Airway submucosal glands are essential for mucus secretion, hydration, and mucociliary clearance, yet the identity and organization of glandular progenitor populations remain incompletely defined. Progress in this area has been limited by the lack of experimental models that recapitulate the gland-rich architecture of human airways. Using the ferret, a species with abundant SMGs throughout the cartilaginous airways, we identify KRT7 as a marker of a conserved glandular ductal stem/progenitor compartment and establish a lineage framework linking airway gland development, maintenance, and regeneration.

A central finding of this study is that KRT7 expression undergoes a developmental transition from broad epithelial expression within the SAE and early stages of SMG morphogenesis to restricted expression within the adult glandular duct epithelium. Unlike humans, ferrets are born with a relatively simple columnar proximal airway epithelium that lacks ciliated and secretory cells and instead contains immature progenitors that give rise to the major airway epithelial lineages during postnatal development ^40,61,64^. Thus, the developmental restriction of KRT7 expression suggests that it marks a progenitor state rather than a lineage-restricted cell population. In the adult airway, the KRT7^+^ ductal compartment overlaps with established progenitor-associated airway basal and gland ductal markers ^46^, including KRT19, CD166, and ALDH1A1, and is conserved between ferret and human airways. Consistent with this interpretation, lineage tracing demonstrated that KRT7^+^ cells give rise to both serous and mucous glandular lineages while progressively losing KRT7 expression as differentiation proceeds. Together, these findings identify KRT7 as a marker of a conserved gland ductal progenitor compartment that contributes to maturation and maintenance of the secretory SMG epithelium.

Although KRT7 is transiently co-expressed with KRT5 and KRT14 during early gland development and lineage tracing revealed that KRT7-lineage cells contribute to KRT5⁺ gland ductal basal cells during SMG morphogenesis, these cells contributed minimally to the basal compartment during repair of the adult surface airway epithelium. This restriction is consistent with the adult SMG duct at homeostasis, where KRT7 expression is confined to luminal ductal cells and excluded from the adjacent KRT5^+^ basal cell layer. Notably, isolated KRT7-lineage cells acquired a KRT5^+^ basal-like phenotype in organoid and ALI cultures, indicating a degree of phenotypic plasticity under in vitro conditions. Together, these findings suggest that the differentiation potential of KRT7-lineage cells is constrained by the native adult airway niche and may be broadened in culture.

This behavior is consistent with regenerative paradigms described in the prostate and salivary glands, where progenitor populations remain largely lineage restricted during homeostasis but can exhibit plasticity following injury or experimental perturbation ^65,66^. In the adult salivary gland, distinct basal, myoepithelial, ductal, and acinar progenitor compartments maintain largely lineage-restricted fates during homeostasis and following mild injury ^66^. However, severe injury can induce cellular plasticity, enabling ductal cells to acquire regenerative functions outside their normal lineage boundaries. Similarly, studies of the prostate demonstrate that lineage restriction can be strongly influenced by tissue context. Although basal and luminal epithelial populations are largely self-sustaining in the adult prostate gland, disruption of normal epithelial architecture can reveal latent multipotent progenitor potential ^65^. Notably, lineage-traced luminal prostate progenitors generate organoids containing both luminal and basal cell populations in vitro ^67^, demonstrating that differentiated epithelial identities can be broadened outside their native niche. Together, these studies support the concept that ductal epithelial progenitor behavior is not fixed but is actively constrained by microenvironmental cues and cell-cell interactions, a model that may similarly explain the acquisition of a KRT5^+^ basal-like phenotype by KRT7-lineage airway gland ductal cells in organoid and ALI cultures.

The regional restriction of ALDH1A1 expression within the KRT7^+^ compartment further indicates that the ductal niche is heterogeneous and likely contains multiple progenitor states. Our in vitro studies further support the progenitor properties of KRT7-lineage cells. KRT7-lineage cells exhibited enhanced proliferative capacity in competitive organoid cultures and preferentially generated large organoid structures relative to non-lineage SMG cells. This growth advantage was influenced by pharmacologic modulation of ALDH activity, consistent with previous studies linking ALDH signaling to airway progenitor function ^46^. However, ALDH inhibition did not abolish organoid formation, suggesting that ALDH contributes to, but does not solely govern, expansion of the KRT7-lineage compartment. These findings indicate that KRT7-lineage cells share conserved metabolic features associated with progenitor activity while retaining additional mechanisms that support growth and maintenance.

Beyond their proliferative capacity, KRT7-lineage cells demonstrated broad epithelial differentiation potential. In both organoid and air-liquid interface cultures, lineage-derived cells generated basal, ciliated, goblet, and secretory cell populations and formed mature airway epithelia. KRT7-lineage organoids also exhibited enhanced forskolin-induced swelling relative to non-lineage controls. Although CFTR dependence was not directly tested in the organoid assay, forskolin-induced swelling is commonly associated with CFTR-mediated fluid transport in epithelial organoid systems. In contrast, CFTR-dependent short-circuit current measurements in differentiated ALI cultures were comparable between KRT7-lineage and non-lineage epithelia. This apparent discrepancy may reflect differences in epithelial architecture, lumen formation, or fluid transport properties captured by the organoid assay but not by transepithelial electrophysiologic measurements. Together, these findings demonstrate that KRT7-lineage cells generate functionally mature airway epithelia capable of CFTR-mediated ion transport and robust agonist-stimulated fluid secretion.

This study has several limitations. Although KRT7-lineage tracing identifies a major progenitor population within the glandular ducts, additional glandular progenitors such as myoepithelial cells likely also contribute to airway maintenance and repair ^25,26^. Furthermore, while the ferret closely models human airway gland biology, species-specific differences in niche regulation cannot be excluded. The present study focused on developmental maturation and acute injury; the behavior of KRT7-lineage cells in chronic inflammatory diseases remains unknown. Likewise, the molecular mechanisms linking ALDH activity to progenitor function, as well as the contribution of the surrounding mesenchymal niche, require further investigation. Future studies incorporating single-cell transcriptomic approaches will be important for resolving heterogeneity within the KRT7^+^ compartment and defining the signaling pathways that regulate glandular progenitor behavior.

In summary, this study identifies KRT7 as a conserved marker of a glandular ductal stem/progenitor compartment in the ferret airway. KRT7-lineage cells contribute to gland development, survive severe epithelial injury, regenerate the airway surface, and generate differentiated epithelia capable of CFTR-mediated transport. These findings establish the glandular duct as an important progenitor niche in the airway and provide a framework for understanding epithelial regeneration in gland-rich mammalian airways.

Supplemental text and figures for this article include experimental details, and detailed materials, and figures that further support the key findings presented in the main text.

## Acknowledgments

This work was supported by the National Institute of Health HL165404 (to JFE) and P30 DK072482 (to BD); NHLBI Contract 75N92025C00007 (to JFE); the Cystic Fibrosis Foundation ENGELH21XX0 (to JFE).

## Ethics approval statement

This study was conducted in accordance with protocols approved by the Institutional Animal Care and Use Committees (IACUC) of The University of Iowa (Protocol #1071945) and the University of Alabama at Birmingham (Protocol #23198), in compliance with National Institutes of Health (NIH) standards. Humane/euthanasia endpoints: 25% loss of body weight from the start of the experiments.

## Author Contributions

Z.H. contributed to the experiments, data analysis, and manuscript drafting. C.W., Y.M.,Y.Y., L.Y., and L.J. co-contributed to the experiments and data analysis. Z.Y., X.S., X.L., and J.F.E. contributed to the study design. X.S., X.L., and J.F.E. supervised the study and provided important intellectual content. X.L., and J.F.E. contributed to the critical revision of the manuscript. All authors read and approved the final manuscript.

## Data availability statement

The data generated during this study are available in the published article and its supplementary files; additional data are available from the corresponding author upon reasonable request.

## References

1. Widdicombe JH. Early studies of airway submucosal glands. Am J Physiol Lung Cell Mol Physiol. Jun 1 2019;316(6):L990–L998. doi:10.1152/ajplung.00068.2019

2. Yu W, Moninger TO, Thurman AL, et al. Cellular and molecular architecture of submucosal glands in wild-type and cystic fibrosis pigs. Proceedings of the National Academy of Sciences. 2022;119(4):e2119759119. doi:doi:10.1073/pnas.2119759119

3. Ostedgaard LS, Price MP, Whitworth KM, et al. Lack of airway submucosal glands impairs respiratory host defenses. Elife. Oct 7 2020;9doi:10.7554/eLife.59653

4. Wine JJ, Joo NS. Submucosal glands and airway defense. Proc Am Thorac Soc. 2004;1(1):47–53. doi:10.1513/pats.2306015

5. Yu F, Zhao X, Li C, et al. Airway stem cells: review of potential impact on understanding of upper airway diseases. Laryngoscope. Jul 2012;122(7):1463–9. doi:10.1002/lary.23320

6. Lu S, Liu H, Farley JM, Sr. Macrolide antibiotics inhibit mucus secretion and calcium entry in Swine airway submucosal mucous gland cells. J Pharmacol Exp Ther. Jan 2011;336(1):178–87. doi:10.1124/jpet.110.171561

7. Cho HJ, Joo NS, Wine JJ. Mucus secretion from individual submucosal glands of the ferret trachea. Am J Physiol Lung Cell Mol Physiol. Jul 2010;299(1):L124–36. doi:10.1152/ajplung.00049.2010

8. Ballard ST, Inglis SK. Liquid secretion properties of airway submucosal glands. J Physiol. Apr 1 2004;556(Pt 1):1–10. doi:10.1113/jphysiol.2003.052779

9. Shah BK, Singh B, Wang Y, Xie S, Wang C. Mucus Hypersecretion in Chronic Obstructive Pulmonary Disease and Its Treatment. Mediators Inflamm. 2023;2023:8840594. doi:10.1155/2023/8840594

10. Abrami M, Biasin A, Tescione F, et al. Mucus Structure, Viscoelastic Properties, and Composition in Chronic Respiratory Diseases. Int J Mol Sci. Feb 5 2024;25(3)doi:10.3390/ijms25031933

11. Rogers DF. Airway mucus hypersecretion in asthma: an undervalued pathology? Curr Opin Pharmacol. Jun 2004;4(3):241–50. doi:10.1016/j.coph.2004.01.011

12. May AJ, Teshima THN, Noble A, Tucker AS. FGF10 is an essential regulator of tracheal submucosal gland morphogenesis. Developmental Biology. 2019/07/15/ 2019;451(2):158–166. 10.1016/j.ydbio.2019.03.017

13. Rose MC, Voynow JA. Respiratory tract mucin genes and mucin glycoproteins in health and disease. Physiol Rev. Jan 2006;86(1):245–78. doi:10.1152/physrev.00010.2005

14. Quinton PM. Cystic fibrosis: impaired bicarbonate secretion and mucoviscidosis. Lancet. Aug 2 2008;372(9636):415–7. doi:10.1016/S0140-6736(08)61162-9

15. Schulz BL, Sloane AJ, Robinson LJ, et al. Mucin glycosylation changes in cystic fibrosis lung disease are not manifest in submucosal gland secretions. Biochem J. May 1 2005;387(Pt 3):911–9. doi:10.1042/bj20041641

16. Meyerholz DK, Stoltz DA, Namati E, et al. Loss of cystic fibrosis transmembrane conductance regulator function produces abnormalities in tracheal development in neonatal pigs and young children. Am J Respir Crit Care Med. Nov 15 2010;182(10):1251–61. doi:10.1164/rccm.201004-0643OC

17. Widdicombe JH. Why Airway Gland Secretions Are Abnormally Sticky in Cystic Fibrosis, and Why Not Much Can Be Done About It. Dev Cell. Aug 24 2020;54(4):427–428. doi:10.1016/j.devcel.2020.07.008

18. Wu JV, Krouse ME, Wine JJ. Acinar origin of CFTR-dependent airway submucosal gland fluid secretion. Am J Physiol Lung Cell Mol Physiol. Jan 2007;292(1):L304–11. doi:10.1152/ajplung.00286.2006

19. Johnson DC. Airway mucus function and dysfunction. N Engl J Med. Mar 10 2011;364(10):978; author reply 978. doi:10.1056/NEJMc1014719

20. Engelhardt JF, Schlossberg H, Yankaskas JR, Dudus L. Progenitor cells of the adult human airway involved in submucosal gland development. Development. Jul 1995;121(7):2031–46. doi:10.1242/dev.121.7.2031

21. Dakir EH, Feigenbaum L, Linnoila RI. Constitutive expression of human keratin 14 gene in mouse lung induces premalignant lesions and squamous differentiation. Carcinogenesis. Dec 2008;29(12):2377–84. doi:10.1093/carcin/bgn190

22. Ievlev V, Jensen-Cody CC, Lynch TJ, et al. Sox9 and Lef1 Regulate the Fate and Behavior of Airway Glandular Progenitors in Response to Injury. Stem Cells. Aug 25 2022;40(8):778–790. doi:10.1093/stmcls/sxac038

23. Borthwick DW, Shahbazian M, Krantz QT, Dorin JR, Randell SH. Evidence for stem-cell niches in the tracheal epithelium. Am J Respir Cell Mol Biol. Jun 2001;24(6):662–70. doi:10.1165/ajrcmb.24.6.4217

24. Hegab AE, Ha VL, Gilbert JL, et al. Novel stem/progenitor cell population from murine tracheal submucosal gland ducts with multipotent regenerative potential. Stem Cells. Aug 2011;29(8):1283–93. doi:10.1002/stem.680

25. Lynch TJ, Anderson PJ, Rotti PG, et al. Submucosal Gland Myoepithelial Cells Are Reserve Stem Cells That Can Regenerate Mouse Tracheal Epithelium. Cell Stem Cell. May 3 2018;22(5):653–667.e5. doi:10.1016/j.stem.2018.03.017

26. Tata A, Kobayashi Y, Chow RD, et al. Myoepithelial Cells of Submucosal Glands Can Function as Reserve Stem Cells to Regenerate Airways after Injury. Cell Stem Cell. May 3 2018;22(5):668–683 e6. doi:10.1016/j.stem.2018.03.018

27. Anderson PJ, Lynch TJ, Engelhardt JF. Multipotent Myoepithelial Progenitor Cells Are Born Early during Airway Submucosal Gland Development. Am J Respir Cell Mol Biol. Jun 2017;56(6):716–726. doi:10.1165/rcmb.2016-0304OC

28. Hegab AE, Ha VL, Attiga YS, Nickerson DW, Gomperts BN. Isolation of basal cells and submucosal gland duct cells from mouse trachea. J Vis Exp. Sep 14 2012;(67):e3731. doi:10.3791/3731

29. Lynch TJ, Engelhardt JF. Progenitor cells in proximal airway epithelial development and regeneration. J Cell Biochem. Oct 2014;115(10):1637–45. doi:10.1002/jcb.24834

30. Di YP, Mou H. Airway Serous Cells: A Comparative Study of Spatial Distribution and Abundance among Species. J Respir Biol Transl Med. Sep 2024;1(3)doi:10.35534/jrbtm.2024.10013

31. Choi HK, Finkbeiner WE, Widdicombe JH. A comparative study of mammalian tracheal mucous glands. J Anat. Oct 2000;197 Pt 3(Pt 3):361–72. doi:10.1046/j.1469-7580.2000.19730361.x

32. Grubb BR, Boucher RC. Pathophysiology of gene-targeted mouse models for cystic fibrosis. Physiol Rev. Jan 1999;79(1 Suppl):S193–214. doi:10.1152/physrev.1999.79.1.S193

33. Widdicombe JH, Wine JJ. Airway Gland Structure and Function. Physiol Rev. Oct 2015;95(4):1241–319. doi:10.1152/physrev.00039.2014

34. Wilke M, Buijs-Offerman RM, Aarbiou J, et al. Mouse models of cystic fibrosis: phenotypic analysis and research applications. J Cyst Fibros. Jun 2011;10 Suppl 2:S152–71. doi:10.1016/s1569-1993(11)60020-9

35. Lee RJ, Foskett JK. Why mouse airway submucosal gland serous cells do not secrete fluid in response to cAMP stimulation. J Biol Chem. Nov 2 2012;287(45):38316–26. doi:10.1074/jbc.M112.412817

36. Raju SV, Kim H, Byzek SA, et al. A ferret model of COPD-related chronic bronchitis. JCI Insight. Sep 22 2016;1(15):e87536. doi:10.1172/jci.insight.87536

37. Smith RE, Choudhary S, Ramirez JA. Ferrets as Models for Viral Respiratory Disease. Comp Med. Jun 1 2023;73(3):187–193. doi:10.30802/aalas-cm-22-000064

38. Wu S, Driver I, Luo M, et al. Ferret model of bleomycin-induced lung injury shares features of human idiopathic pulmonary fibrosis. NPJ Regen Med. Nov 21 2025;10(1):53. doi:10.1038/s41536-025-00440-z

39. Yan Z, Stewart ZA, Sinn PL, et al. Ferret and pig models of cystic fibrosis: prospects and promise for gene therapy. Hum Gene Ther Clin Dev. Mar 2015;26(1):38–49. doi:10.1089/humc.2014.154

40. Sehgal A, Presente A, Engelhardt JF. Developmental expression patterns of CFTR in ferret tracheal surface airway and submucosal gland epithelia. Am J Respir Cell Mol Biol. Jul 1996;15(1):122–31. doi:10.1165/ajrcmb.15.1.8679216

41. Keiser NW, Birket SE, Evans IA, et al. Defective innate immunity and hyperinflammation in newborn cystic fibrosis transmembrane conductance regulator-knockout ferret lungs. Am J Respir Cell Mol Biol. Jun 2015;52(6):683–94. doi:10.1165/rcmb.2014-0250OC

42. Rosen BH, Evans TIA, Moll SR, et al. Infection Is Not Required for Mucoinflammatory Lung Disease in CFTR-Knockout Ferrets. Am J Respir Crit Care Med. May 15 2018;197(10):1308–1318. doi:10.1164/rccm.201708-1616OC

43. Sun X, Sui H, Fisher JT, et al. Disease phenotype of a ferret CFTR-knockout model of cystic fibrosis. J Clin Invest. Sep 2010;120(9):3149–60. doi:10.1172/jci43052

44. Hegab AE, Ha VL, Darmawan DO, et al. Isolation and in vitro characterization of basal and submucosal gland duct stem/progenitor cells from human proximal airways. Stem Cells Transl Med. Oct 2012;1(10):719–24. doi:10.5966/sctm.2012-0056

45. Liu X, Engelhardt JF. The glandular stem/progenitor cell niche in airway development and repair. Proc Am Thorac Soc. Aug 15 2008;5(6):682–8. doi:10.1513/pats.200801-003AW

46. Hegab AE, Ha VL, Bisht B, et al. Aldehyde dehydrogenase activity enriches for proximal airway basal stem cells and promotes their proliferation. Stem Cells Dev. Mar 15 2014;23(6):664–75. doi:10.1089/scd.2013.0295

47. Sun X, Yi Y, Xie W, et al. CFTR Influences Beta Cell Function and Insulin Secretion Through Non-Cell Autonomous Exocrine-Derived Factors. Endocrinology. Oct 1 2017;158(10):3325–3338. doi:10.1210/en.2017-00187

48. Yu M, Sun X, Tyler SR, et al. Highly Efficient Transgenesis in Ferrets Using CRISPR/Cas9-Mediated Homology-Independent Insertion at the ROSA26 Locus. Sci Rep. Feb 13 2019;9(1):1971. doi:10.1038/s41598-018-37192-4

49. He N, Liu X, Vegter AR, et al. Ferret models of alpha-1 antitrypsin deficiency develop lung and liver disease. JCI Insight. Mar 8 2022;7(5)doi:10.1172/jci.insight.143004

50. Yuan F, Gasser GN, Lemire E, et al. Transgenic ferret models define pulmonary ionocyte diversity and function. Nature. Sep 2023;621(7980):857–867. doi:10.1038/s41586-023-06549-9

51. Ievlev V, Pai AC, Dillon DS, Jr., et al. Development and characterization of ferret ex vivo tracheal injury and cell engraftment model. Front Med (Lausanne). 2023;10:1144754. doi:10.3389/fmed.2023.1144754

52. Dekkers JF, Berkers G, Kruisselbrink E, et al. Characterizing responses to CFTR-modulating drugs using rectal organoids derived from subjects with cystic fibrosis. Sci Transl Med. Jun 22 2016;8(344):344ra84. doi:10.1126/scitranslmed.aad8278

53. Yang L, Yao C, Hajibabazade J, et al. CFTR facilitates fluid secretion by ferret alveolar type 2 cells. J Cyst Fibros. May 15 2026;doi:10.1016/j.jcf.2026.05.004

54. Yuan F, Hajibabazade J, Yi Y, et al. Ionocyte CFTR Coordinates Chloride Absorption and Secretion to Balance Airway Fluid. Am J Respir Crit Care Med. Aug 15 2025;doi:10.1164/rccm.202505-1242OC

55. Liu X, Luo M, Zhang L, Ding W, Yan Z, Engelhardt JF. Bioelectric properties of chloride channels in human, pig, ferret, and mouse airway epithelia. Am J Respir Cell Mol Biol. Mar 2007;36(3):313–23. doi:10.1165/rcmb.2006-0286OC

56. Hajighasemi-Ossareh M, Borthwell RM, Lachowicz-Scroggins M, Stevens JE, Finkbeiner WE, Widdicombe JH. Distribution and size of mucous glands in the ferret tracheobronchial tree. Anat Rec (Hoboken). Nov 2013;296(11):1768–74. doi:10.1002/ar.22783

57. Alam CM, Baghestani S, Pajari A, Omary MB, Toivola DM. Keratin 7 Is a Constituent of the Keratin Network in Mouse Pancreatic Islets and Is Upregulated in Experimental Diabetes. Int J Mol Sci. Jul 21 2021;22(15)doi:10.3390/ijms22157784

58. Hosseinalizadeh H, Hussain QM, Poshtchaman Z, et al. Emerging insights into keratin 7 roles in tumor progression and metastasis of cancers. Front Oncol. 2023;13:1243871. doi:10.3389/fonc.2023.1243871

59. Hrudka J, Fiserova H, Jelinkova K, Matej R, Waldauf P. Cytokeratin 7 expression as a predictor of an unfavorable prognosis in colorectal carcinoma. Sci Rep. Sep 9 2021;11(1):17863. doi:10.1038/s41598-021-97480-4

60. Jiang M, Li H, Zhang Y, et al. Transitional basal cells at the squamous-columnar junction generate Barrett’s oesophagus. Nature. Oct 26 2017;550(7677):529–533. doi:10.1038/nature24269

61. Leigh MW, Gambling TM, Carson JL, Collier AM, Wood RE, Boat TF. Postnatal development of tracheal surface epithelium and submucosal glands in the ferret. Exp Lung Res. 1986;10(2):153–69. doi:10.3109/01902148609061490

62. Duan D, Yue Y, Zhou W, et al. Submucosal gland development in the airway is controlled by lymphoid enhancer binding factor 1 (LEF1). Development. Oct 1999;126(20):4441–53. doi:10.1242/dev.126.20.4441

63. Hegab AE, Nickerson DW, Ha VL, Darmawan DO, Gomperts BN. Repair and regeneration of tracheal surface epithelium and submucosal glands in a mouse model of hypoxic-ischemic injury. Respirology. Oct 2012;17(7):1101–13. doi:10.1111/j.1440-1843.2012.02204.x

64. Voter KZ, Leigh MW, Boat TF, Carson JL, Wood RE. Development of mucociliary transport in the postnatal ferret trachea. J Appl Physiol (1985). Oct 1992;73(4):1500-3. doi:10.1152/jappl.1992.73.4.1500

65. Horton C, Liu Y, Yu C, Xie Q, Wang ZA. Luminal-contact-inhibition of epithelial basal stem cell multipotency in prostate organogenesis and homeostasis. Biol Open. Oct 1 2019;8(10)doi:10.1242/bio.045724

66. Ninche N, Kwak M, Ghazizadeh S. Diverse epithelial cell populations contribute to the regeneration of secretory units in injured salivary glands. Development. Oct 9 2020;147(19)doi:10.1242/dev.192807

67. Chua CW, Shibata M, Lei M, et al. Single luminal epithelial progenitors can generate prostate organoids in culture. Nat Cell Biol. Oct 2014;16(10):951–61, 1-4. doi:10.1038/ncb3047

