## Supplemental data for "KRT7 Marks an Airway Glandular Ductal Stem Cell Population That Supports Multilineage Development and Regeneration"

**Running head: KRT7**[+] Airway Glandular Stem Cells

**Acknowledgments:** This work was supported by the National Institute of Health HL165404 (to JFE) and P30 DK072482 (to BD); NHLBI Contract 75N92025C00007 (to JFE); the Cystic Fibrosis Foundation ENGELH21XX0 (to JFE).

**Online Supplemental Data**

**Supplemental Materials and Methods**

**Supplemental Figures**

**Supplemental Tables**

### Supplemental Materials and Methods

#### *Ferrets*

The Institutional Animal Care and Use Committee of the University of Alabama at Birmingham approved the animal experimentation. Male and female domestic ferrets aged 3 to 24 months were purchased from Marshall Farms in North Rose, NY, USA, while kit ferrets were bred at campus animal facilities. The ferrets were housed at a controlled temperature of 20–22°C under a 16-hour light and 8-hour dark cycle, with unrestricted access to water and food. Animals were used for experiments after a two-week acclimatization period. Selection was based on genotype, and blinding was unnecessary because the quantitative detection method was unbiased. For transgenic ferrets, sex was not considered a variable owing to the difficulty in evenly distributing sexes among them.

#### *Lineage Tracing of KRT7-lineage Cells in $KRT7Cre^{ERT2}$:: $ROSA^{mTmG}$ Ferrets*

To perform lineage tracing of airway glandular stem cells (GDSCs), $KRT7^{CreERT2}$ transgenic ferrets were crossed with ROSA-TG Cre-reporter ferrets to generate hemizygous offspring (Yu *et al.*, 2019). The ferrets received intraperitoneal injections of tamoxifen (20 mg/kg in sunflower seed oil) daily for five consecutive days to induce Cre-mediated recombination. The animals were euthanized for lineage tracing analysis at predetermined experimental timepoints or underwent intrapulmonary injury with polidocanol two weeks after tamoxifen induction.

#### *Polidocanol injury of ferret airways*

Following tamoxifen-induced genetic labeling of KRT7-expressing cells, ferrets were subjected to injury through the administration of a single dose of polidocanol (2%, ml/kg in saline, Thesit, Cas No. 9002-92-0, Sigma, St. Louis, MO) via the laryngotracheal route utilizing a MADgic® Laryngo-Tracheal Mucosal Atomization Device (Teleflex, Morrisville, NC) two weeks after the final dose of tamoxifen. All ferrets were monitored daily for activity level, body weight, and respiratory rate during the initial week following polidocanol administration. Humane endpoints included: 1) hemoptysis three days post-injury;

2) loss of more than 20% of body weight at any point during the first week; 3) labored respiration with a respiratory rate exceeding 90 breaths per minute; 4) lack of movement or response to stimulation. Animals were euthanized at predetermined experimental timepoints for analysis or reached a humane endpoint.

***Colocalization Analysis of KRT7-Lineage Cells with Epithelial Markers***

Colocalization analysis was performed in Fiji to quantify overlap between $GFP^+$ KRT7-lineage cells and specific epithelial or glandular cell markers in immunofluorescence images. Multichannel fluorescence images were first split into individual channels, including DAPI, GFP, and marker-specific channels. Nuclei were identified from DAPI staining using threshold-based segmentation, and individual cells were defined as regions of interest (ROIs) using the "Analyze Particles" function, with size and circularity filters applied to exclude debris and overlapping nuclei. For each marker channel, fluorescence was thresholded to distinguish positive signal from background. The mean fluorescence intensity within each DAPI-defined ROI was measured, and a cutoff for marker positivity was set based on the distribution of intensity values across all cells in the image. Cells with a mean intensity above the threshold were classified as marker positive. $GFP^+$ KRT7-lineage cells were identified using the same thresholding and ROI-based approach. Colocalization was assessed at the single-cell level by identifying ROIs positive for both GFP and the corresponding marker. Cells meeting both criteria within the same ROI were defined as double positive. The proportion of colocalized cells was calculated relative to the total number of $DAPI^+$ cells or $GFP^+$ lineage cells, depending on the specific analysis. All threshold parameters were kept consistent across images within each experimental group to ensure unbiased quantification. Multiple fields per sample were analyzed, and data from independent biological replicates were pooled for statistical analysis.

***Organoid Culture and Treatment***

FACS-sorted GFP-positive KRT7-lineage cells and tdTomato-positive non-lineage cells were mixed at a 1:1 ratio and embedded in growth factor-reduced Matrigel (Cat# 354230, Corning, Glendale, AZ). For each well of a 24-well plate, 1,000 GFP-positive cells and 1,000 tdTomato-positive cells were suspended in a total volume of 50 µL Matrigel and plated as a dome. After gel polymerization at 37°C for 15–30 min,

500 μL of PneumaCult Ex-Plus medium (Stemcell Technologies) was added to each well. Organoids were maintained without treatment from day 0 to day 3 to allow initial establishment.

On day 3, brightfield and fluorescence images were acquired, and the medium was replaced with Ex-Plus medium containing one of the following treatments: DMSO vehicle control, Alda-1, Daidzin, or DEAB. Alda-1 was obtained from MedChemExpress, whereas Daidzin and DEAB were purchased from Sigma-Aldrich (Daidzin, CAS No. 552-66-9; DEAB [4-diethylaminobenzaldehyde], CAS No. 120-21-8). All compounds were used at a final concentration of 20 μM. Drug-containing medium was prepared by diluting stock solutions in Ex-Plus medium (3 μL stock per 3 mL medium). The medium was refreshed on day 7 under the same treatment conditions. On day 10, organoids were imaged, after which the medium was replaced with PneumaCult ALI medium without drug supplementation. Cultures were then maintained until day 21.

On day 21, organoids from the DMSO control group were collected for downstream analyses. A subset was used for swelling assays, while the remaining organoids were fixed, cryoprotected in 20% sucrose, embedded in OCT, cryosectioned, and processed for immunofluorescence staining.

**References**

Yu M, Sun X, Tyler SR, Liang B, Swatek AM, Lynch TJ, He N, Yuan F, Feng Z, Rotti PG, Choi SH, Shahin W, Liu X, Yan Z, Engelhardt JF (2019). Highly Efficient Transgenesis in Ferrets Using CRISPR/Cas9-Mediated Homology-Independent Insertion at the ROSA26 Locus. Sci Rep 9, 1971.

### Supplemental Figures

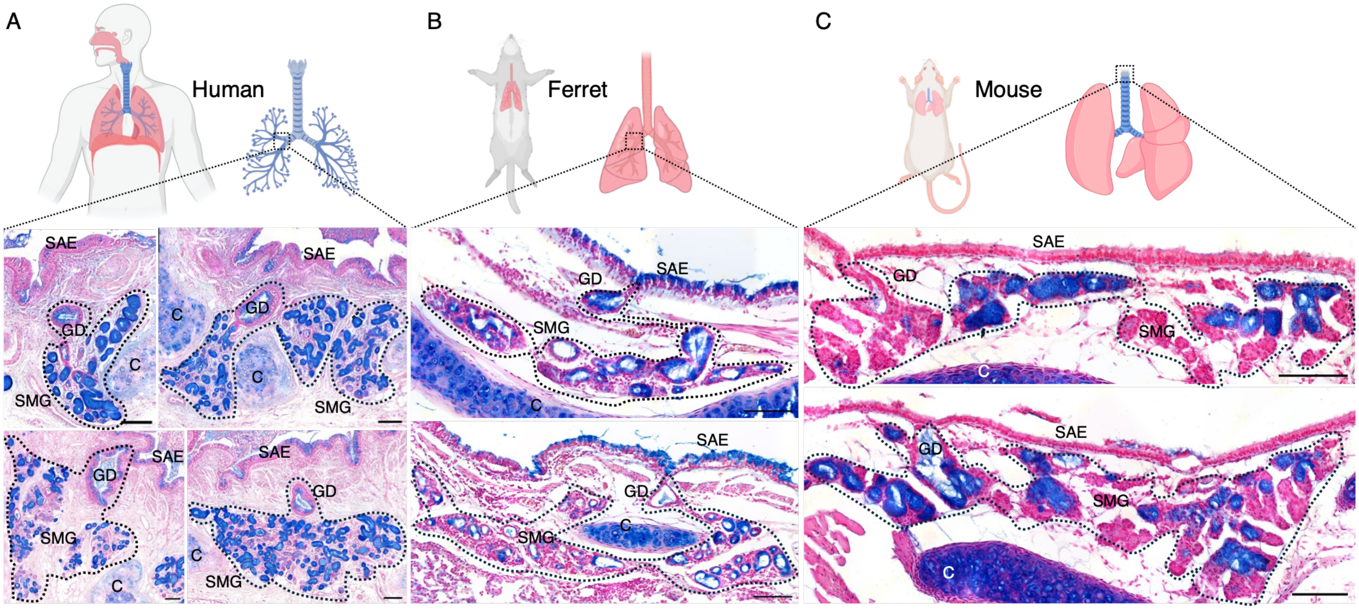


**Supplementary Figure S1. Comparative anatomical and histological distribution of airway submucosal glands across human, ferret, and murine models.** (A-C) Representative histological images of bronchial sections of human (A) and ferret (B), and the proximal trachea of the mouse (C) stained with Alcian Blue to identify mucin-producing structures. Alcian Blue highlights acidic mucins within the surface airway epithelium (SAE) and airway submucosal glands (SMGs). Abundant SMGs are observed in human and ferret bronchus, whereas glandular structures are sparse and are only present in the proximal trachea. Alcian Blue stains acidic mucins (dark blue) in the surface airway epithelium (SAE) and the acinar cells of the airway submucosal glands (SMG), while the pink/red counterstain highlights general cellular and tissue architecture. Both human and ferret bronchi exhibit a thick submucosal layer heavily populated with abundant, well-developed SMGs. In stark contrast, the mouse proximal trachea demonstrates a notably thin submucosal space with cartilage situated immediately beneath the epithelium, and SMGs are sparse and highly restricted to the uppermost proximal trachea. Black dotted lines delineate the distinct anatomical boundaries of the SMG clusters within the submucosa. Scale bars equal 100 µm in all images. SAE, surface airway epithelium; SMG, submucosal gland; GD, gland duct; C, cartilage.

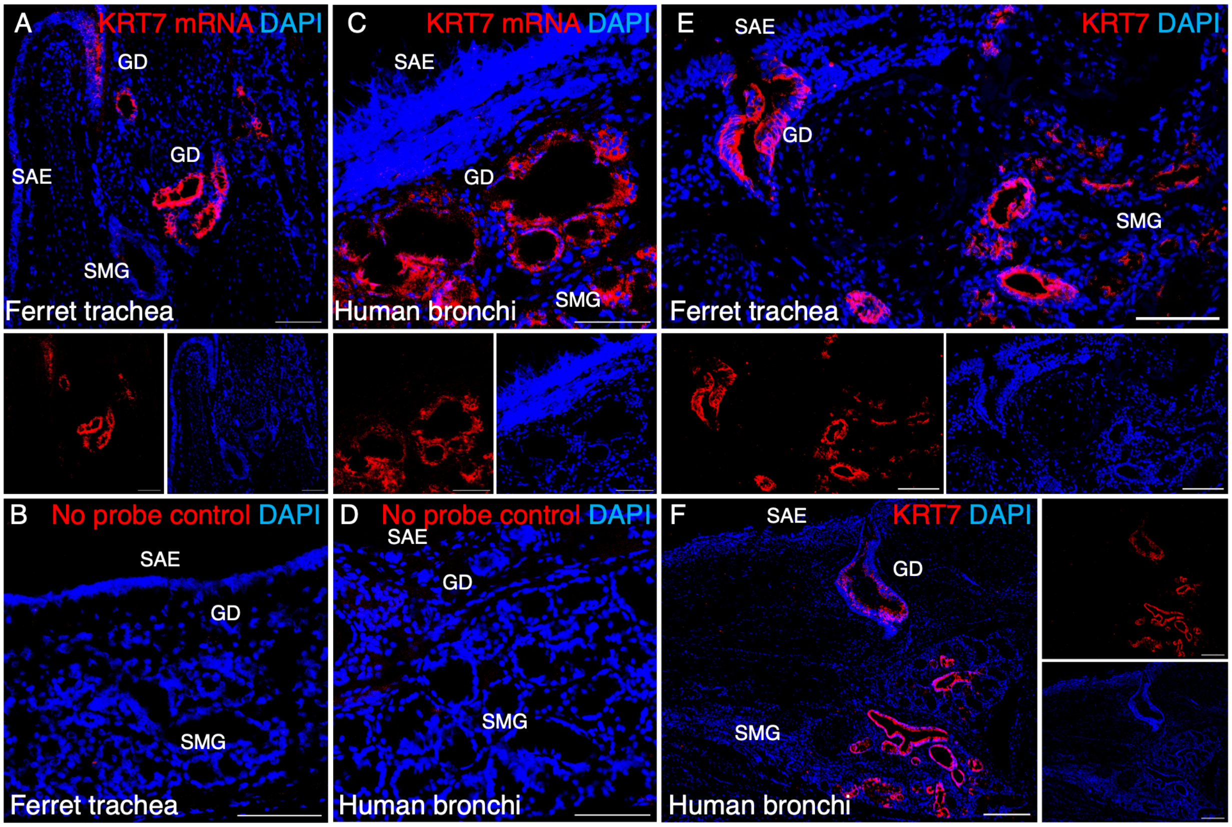


**Supplementary Figure S2. Conserved expression of KRT7 mRNA and protein specifically localizes to the glandular ductal epithelia in ferret and human airways.** (A-D) KRT7 mRNA expression: Representative images of single-molecule fluorescence RNA in situ hybridization (red). Robust KRT7 mRNA expression is highly localized to the epithelial cells of the gland duct (GD) in both the (A) ferret trachea and (C) human bronchi. The lack of signal in the corresponding no-probe controls for both ferret (B) and human (D) tissues confirms the strict specificity of the assay. (E, F) KRT7 protein expression: Representative immunofluorescence images showing that KRT7 protein (red) strongly mirrors the transcriptional mRNA patterns. At the translational level, KRT7 is exclusively concentrated within the GD epithelia of the (E) ferret trachea and (F) human bronchi, with minimal to no expression in the adjacent surface or deep acinar tissues. Lower insets in panels (A, C, E) and the right-side inset in panel (F) display isolated single-channel fluorescence or merged context to clearly delineate the specific KRT7 signal from the DAPI overlay. Nuclei were counterstained with DAPI. Scale bars are 100 µm in all panels. GD, gland duct; SAE, surface airway epithelium; SMG, submucosal gland.

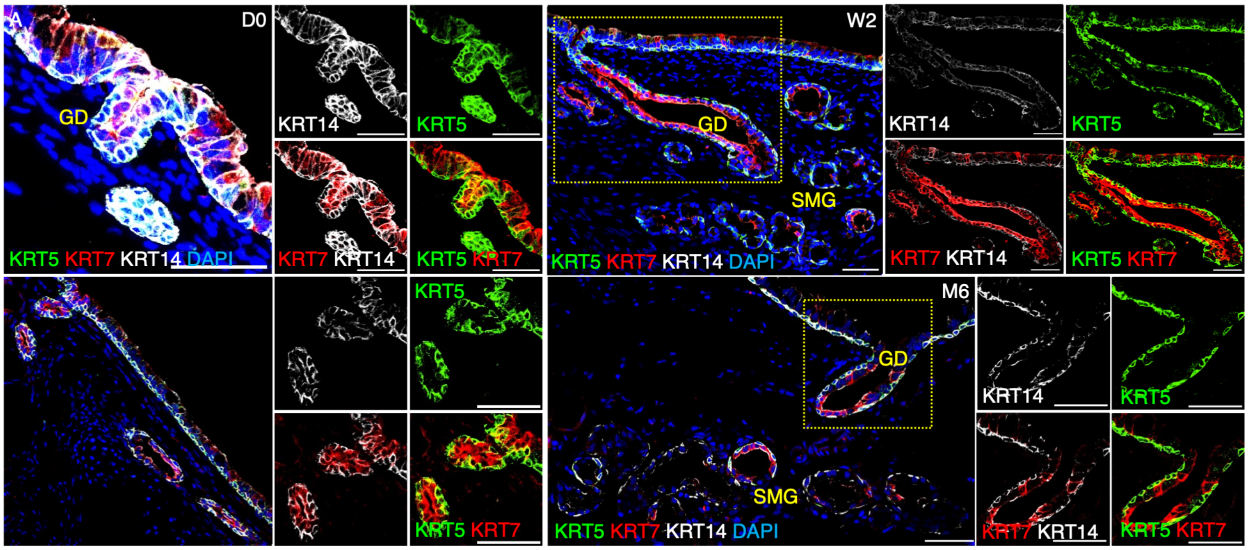


**Supplementary Figure S3. Spatiotemporal mapping of KRT7 relative to basal cell markers KRT5 and KRT14 during ferret airway submucosal gland (SMG) development.** (A-D) Representative immunofluorescence images of ferret tracheal sections across progressive stages of postnatal development: (A) Day 0 (D0, newborn), (B) 2 weeks (W2), (C) 4 weeks (W4), and (D) 6 months (M6, adult). Tissues were co-stained to evaluate the spatial relationship between KRT7 (red) and the established basal cell markers KRT5 (green) and KRT14 (gray/white). During the initial formation of the gland bud (GB), KRT7 was broadly expressed throughout the invaginating epithelium. Extensive co-localization was observed between KRT7 and the basal markers KRT5/KRT14 within these primitive structures. As the glands mature and structurally diversify into distinct ducts (GD) and deep submucosal acini, KRT5 and KRT14 progressively restrict to the basal and myoepithelial layers. Concurrently, KRT7 expression begins to concentrate within developing gland ducts, highlighting the stratified ductal epithelium, where it partially overlaps with basal markers and extends into the luminal layers. In the fully mature adult trachea, expression profiles are highly compartmentalized. While KRT5 and KRT14 continuously define the basal cell layer across the surface airway epithelium (SAE), GD, and SMG, KRT7 expression was strictly restricted to the stratified epithelium of the GD and is excluded from the deep SMG acini. The inset panels on the right side of each main image display the corresponding single- or dual-channel fluorescence, clearly delineating signal overlap and spatial segregation. Nuclei were counterstained with DAPI. Scale bars are 100 µm in all panels. GB, gland bud; GD, gland duct; SAE, surface airway epithelium; SMG, submucosal gland.

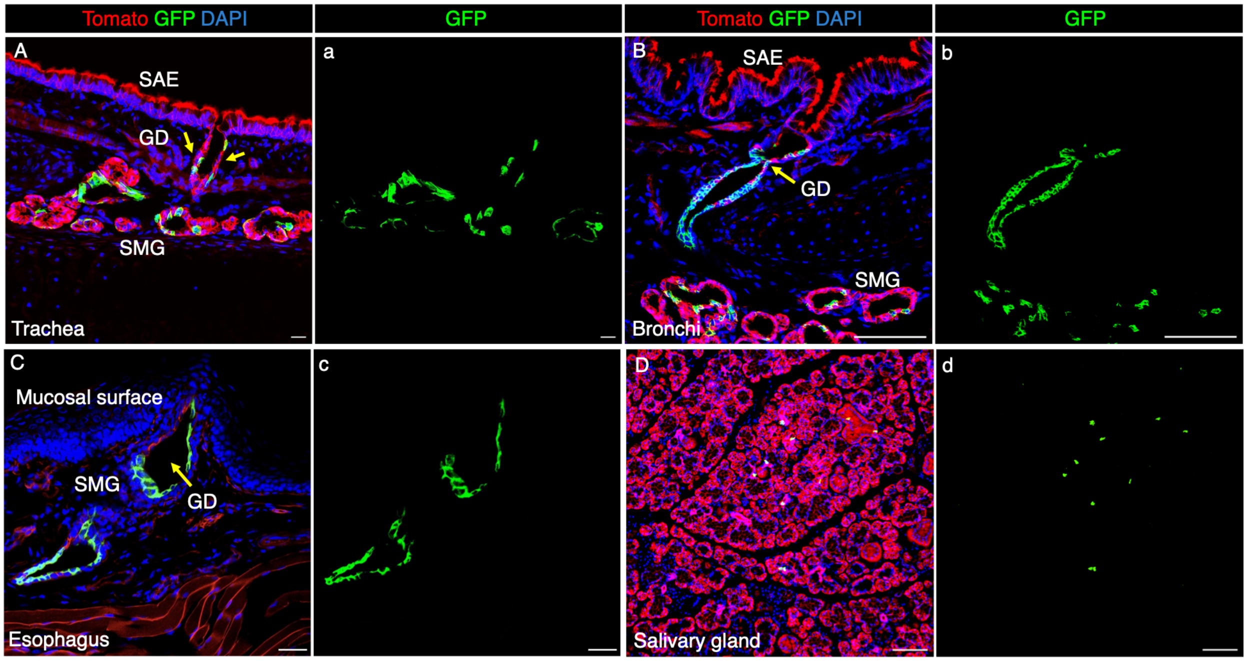


**Supplementary Figure S4. In vivo KRT7-lineage labeling specifically targets glandular ductal epithelial cells across distinct mucosal tissues in the ferret.** (A–D) Representative fluorescence images of the indicated glandular organs from tamoxifen-induced *KRT7*$^{\mathrm{CreER2}}$::*ROSA*$^{\mathrm{mTmG}}$ adult ferrets. Successful recombination permanently labels KRT7-lineage cells with GFP (green), while unrecombined, non-KRT7-lineage cells express tdTomato (red). Nuclei are counterstained with DAPI (blue). In the cartilage airway, robust and highly specific GFP labeling was localized exclusively to the distinct invaginations of the gland ducts (GD, indicated by yellow arrows) of the trachea (A), bronchus (B), and esophagus (C). The surface airway epithelium (SAE) and the deeper submucosal gland (SMG) acini remained Tomato-positive, highlighting the duct-specific nature of the KRT7 lineage. In contrast to the airway, the salivary gland (D) exhibits extremely sparse KRT7-lineage (GFP$^{+}$) labeling, with most of the glandular architecture remaining tdTomato$^{+}$. Although GFP expression in the esophagus was strictly confined to the ductal structures (GD, yellow arrow) of the esophageal SMGs, the KRT7 lineage was excluded from both the deep SMG secretory cells and the thick stratified squamous surface epithelium (SAE). The lowercase panels (a–d) display the corresponding isolated GFP (green) single-channel images, clearly delineating the precise spatial distribution of the KRT7 lineage without the tdTomato signal overlay. Scale bars equal 50 µm in all panels. GD, gland duct; SAE, surface airway epithelium in (A, B), or squamous surface epithelium in (D); SMG, submucosal gland.

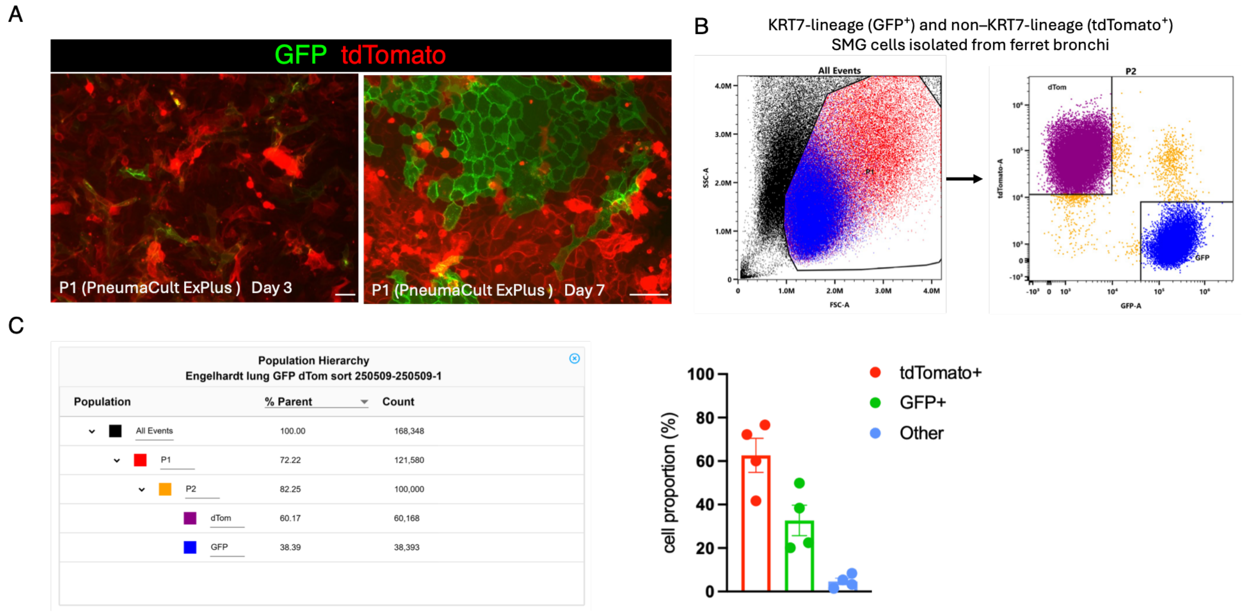


**Supplementary Figure S5. Isolation and separation of GFP$^+$ KRT7-lineage cells and tdTomato$^+$ non-KRT7 lineage cells from bronchial submucosal glands of tamoxifen-induced *KRT7*$^{CreER2}$::*ROSA*$^{mTmG}$ ferrets.** (A) Representative immunofluorescence image of cultured bronchial total SMG cells showing GFP$^+$ KRT7-lineage cells and tdTomato$^+$ non-KRT7 lineage cells in 2-D cultures at passage 0 at Day 3 (left panel) and Day 7 (right panel) in PneumaCult Ex-Plus medium. The image demonstrates distinct, non-overlapping populations of GFP$^+$ KRT7-lineage cells (green) and tdTomato$^+$ non-KRT7-lineage cells (red) that maintain robust reporter fluorescence. Scale bar: 100 µm. (B) Representative fluorescence-activated cell sorting (FACS) plots detailing the separation protocol. The left plot (FSC-A vs. SSC-A) outlines the initial gating for viable cell populations (P1). The right plot shows clear fluorescence separation of the selected viable cells into purified tdTomato$^+$ (tdTomato, purple) and GFP$^+$ (blue) target populations, while excluding double-positive cells and unstained debris. (C) A representative population hierarchy table summarizing the sorting yields. It tracks the step-by-step refinement from total events down to the final sorted tdTomato (approx. 60% of the P2 parent gate) and GFP (approx. 38% of the P2 parent gate) populations, ensuring high purity of the isolated fractions. (D) Bar graph displaying the overall composition of the isolated SMG cell fractions. The data quantify the percentages of tdTomato$^+$ non-KRT7 lineage cells (red bar), GFP$^+$ KRT7-lineage cells (green bar), and unlabeled cells (blue bar) within the total SMG cell prep. Individual data points (N=4) represent values from independent donor ferrets. Data are presented as mean ± SEM. The distinct data points in the bar graph represent the count for each independent experiment.

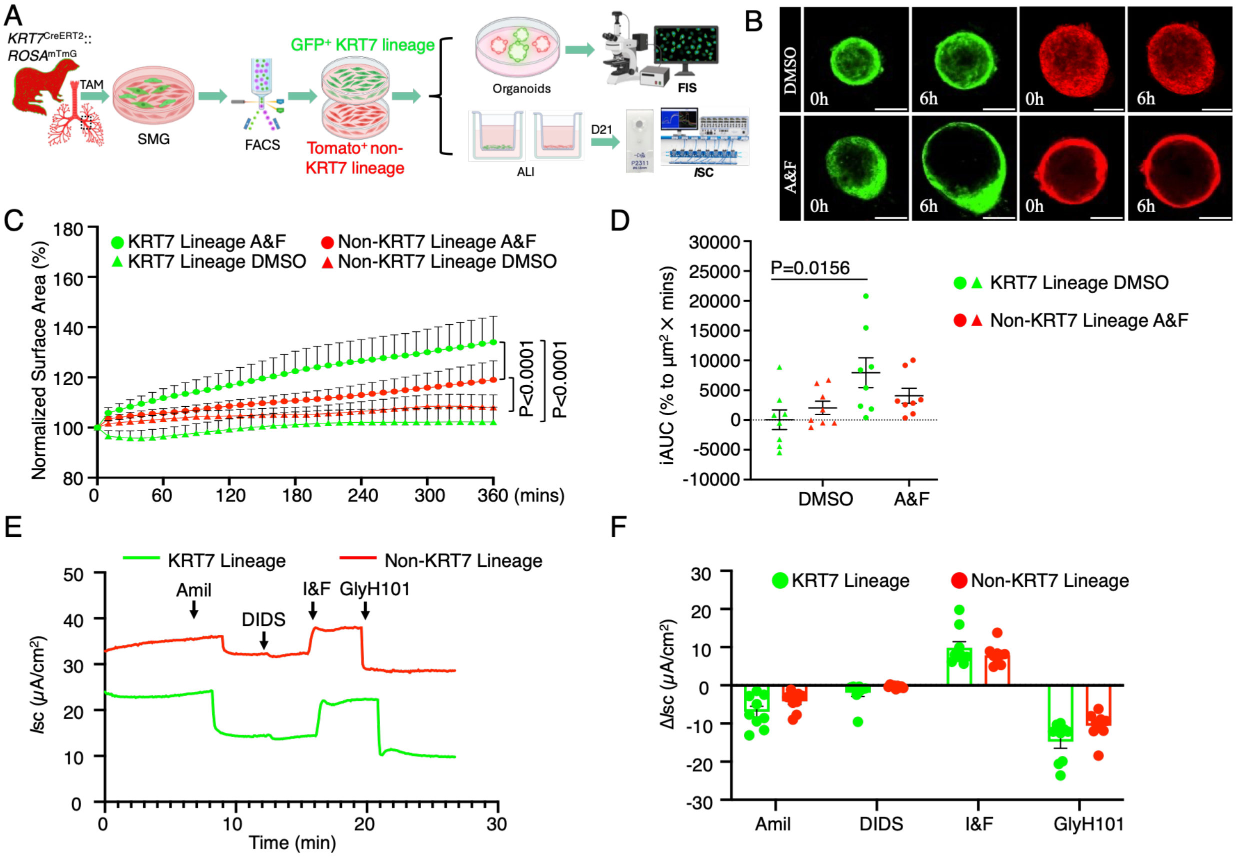


**Supplementary Figure S6. Robust CFTR-mediated ion and fluid transport by epithelia derived from KRT7 glandular ductal stem cell (GDSC) lineages.** (A) Schematic detailing the in vitro culture of organoids and polarized epithelia derived from ferret GFP$^{+}$ KRT7-lineage cells and Tomato$^{+}$ non-KRT7-lineage cells for assessing CFTR-mediated ion transport by forskolin-induced swelling (FIS) assay and Ussing chamber short-circuit current (*I*sc) assay, respectively. Adult *KRT7*$^{\text{CreERT2}}$::*ROSA*$^{\text{mTmG}}$ ferrets were administered tamoxifen for KRT7 lineage labeling. Submucosal gland (SMG) cells were isolated and dissociated, followed by fluorescence-activated cell sorting (FACS) to separate the KRT7 lineage (GFP$^{+}$) from the non-KRT7 lineage (tdTomato$^{+}$). Sorted cells were propagated in PneumaClut Ex+ medium before they were seeded in Matrigel for organoid culture and on Transwell membrane for air-liquid interface (ALI) culture. CFTR-mediated ion transport was measured after 14 days post-differentiation. (B) Representative time-lapse fluorescence images (0 and 6 hrs) of GFP$^{+}$ KRT7 lineage- and tdTomato$^{+}$ non-GDSClineage-derived organoids. Organoids were treated with either vehicle control (DMSO) or a combination of Forskolin and Amiloride (A&F). Forskolin strongly stimulates cAMP-dependent CFTR activation, while Amiloride inhibits ENaC (epithelial sodium channels) to isolate the CFTR-driven chloride and fluid secretion. A pronounced, visible swelling is exclusively observed in the GFP$^{+}$ KRT7-lineage organoids after 6 hours of A&F treatment. (C) Traces showing the real-time changes in total organoid surface area (normalized as a percentage of the baseline area at 0 hours) over a 360-minute (6-hour) period. KRT7-lineage organoids treated with A&F (green circles) exhibited a rapid and significant increase in surface area compared to their DMSO controls (green triangles) and all non-KRT7 lineage groups. (D) Quantification of the integrated area under the curve (iAUC) for GDSC and non-GDSC lineage organoids treated with DMSO or A&F. The $GFP^+$ KRT7-lineage organoids demonstrated significantly higher iAUC upon A&F stimulation compared to their baseline DMSO controls (P=0.0156). N=8 independent organoid cultures derived from 2 donor ferrets. (D) Representative traces of short-circuit current (*I*sc) of polarized epithelia derived KRT7 lineage and non-KRT7 glandular cells showing CFTR-dependent *I*sc responses following stimulation with IBMX/forskolin (I&F) in the presence of amiloride and DIDS, and inhibition by the CFTR inhibitor GlyH101 (GlyH). (E) Quantification of changes in *I*sc (Δ*I*sc) in ALI cultures following treatment with the indicated pharmacological agents shows no significant difference between ALI epithelia derived from KRT7- and non-KRT7-lineage cells (N=9 from 3 independent experiments using 3 donor ferrets). Data are presented as the mean ± SEM. Scale bars = 100 μm in panel (B). Statistical significance was assessed using a two-way ANOVA Tukey Multiple comparison for the kinetic traces (B), and a one-way ANOVA Tukey Multiple comparison test for the iAUC comparisons (D). The distinct data points in the bars of each graph represent the average of triplicate counts of each independent experiment per donor.

### Supplementary Tables

#### Supplementary Table S1. List of primary antibodies information

| Antibody | Vendor | Cat # | Species of Origin | Species Against | Application & Dilution |
|---|---|---|---|---|---|
| KRT7 | Invitrogen | MA1-06315 | Mouse | Human, Mouse, Rat | IF (1:200) |
| GFP | Abcam | ab13970 | Chicken | | IF (1:1000) |
| KRT19 | GeneTex | GTX112826 | Rabbit | Human, Mouse, Dog | IF (1:400) |
| ALCAM (CD166) | Cell Signaling | 10623 | Rabbit | Human, Mouse | IF (1:200) |
| ALDH1A1 | Invitrogen | CL594-60171 | Mouse | Human, Mouse, Rat | IF (1:300) |
| LEF1 | Cell Signaling | 2230 | Rabbit | Human, Mouse, Rat | IF (1:200) |
| MUC5B | Sigma | HPA008246 | Rabbit | Human | IF (1:500) |
| Lysozyme | | | | | |
| SMA | Abcam | ab7817 | Mouse | Human | IF (1:500) |
| KRT5 | OriGene | BP5006 | Guinea pig | Human | IF (1:500) |
| KRT5 | BioLegend | 905504 | Rabbit | Mouse | IF (1:500) |
| FOXJ1 | Abcam | ab235445 | Rabbit | Human, Mouse, Rat | IF (1:500) |
| SCGB1A1 | Millipore | ABS1673 | Goat | Rat | IF (1:1000) |
| MUC5AC | Abcam | ab3649 | Mouse | Human, Pig, Rabbit | IF (1:500) |
| Acetyl- α-Tubulin | Cell Signaling | 5335S | Rabbit | Human, Mouse, Rat | IF (1:500) |

**Supplementary Table S2. List of secondary antibodies information**

| Antibody | Vendor | Cat # | Species of Origin | Application & Dilution |
|---|---|---|---|---|
| Alexa Fluor 488-conjugated AffiniPure Donkey Anti-Chicken IgY++ (IgG) (H+L) | Jackson ImmunoResearch | 703-546-155 | Chicken | IF (1:500) |
| Alexa Fluor 568 Donkey Anti-Rabbit IgG (H+L) | Invitrogen | A10042 | Rabbit | IF (1:500) |
| Alexa Fluor 568 donkey anti-goat IgG (H+L) | Invitrogen | A-11057 | Goat | IF (1:500) |
| Alexa Fluor 647-conjugated AffiniPure F(ab')2 Fragment Donkey Anti-Rabbit IgG (H+L) | Jackson ImmunoResearch | 711-606-152 | Rabbit | IF (1:500) |
| Alexa Fluor 647 donkey anti-mouse IgG (H+L) | Molecular Probes | A31571 | Mouse | IF (1:500) |
| Donkey anti-Goat IgG (H+L) Cross-Adsorbed Secondary Antibody, Alexa Fluor 647 | Invitrogen | A-21447 | Goat | IF (1:500) |
| Alexa Fluor® 647-AffiniPure F(ab')2 Fragment Donkey Anti-Guinea Pig IgG (H+L) | Jackson ImmunoResearch | 706-606-148 | Guinea pig | IF (1:500) |

**Supplementary Table S3. List of primers information**

| Primer | Sequence |
|---|---|
| *GAPDH* | Forward 5'CAACTTTGGCATTGTGGAGG |
| | Reverse 5'CAGTGATGGCATGGACGGTGG |
| | Probe 5'CAGTGGAAGCAGGGATGATG |
| *α-tubulin* | Forward 5'TGCTGCTGTTCACCTTCACC |
| | Reverse 5'CGGTGAACTCCATCTCGTCC |
| | Probe 5' |
| *KRT5* | Forward CCGGGAGTGGATTTGGTTT |
| | Reverse ACCAAAGCCGCCTCCAAAGC |
| | Probe 5'CTGCTCAACATTGCTGCCGACC |
| *MUC5AC* | Forward CGTTGCTGACCTCGATGAG |
| | Reverse CAGTCACCATGAAGCTCGAC |
| | Probe AGCTGACCAAAGGCTCCATCCTG |
| *SCGB1A1* | Forward AGTTACGAGGCTGCACTTG |
| | Reverse GGAGTTTTTAAGACTTTGCCCG |
| | Probe AGGATGTTCTGTGTGGTGTTTGAGGG |
